# A quantitative map of human Condensins provides new insights into mitotic chromosome architecture

**DOI:** 10.1101/237834

**Authors:** Nike Walther, M. Julius Hossain, Antonio Z. Politi, Birgit Koch, Moritz Kueblbeck, Øyvind Ødegård-Fougner, Marko Lampe, Jan Ellenberg

**Author notes:** Correspondence (J.E.). Max Planck Institute for Medical Research, 69120 Heidelberg, Germany (Birgit Koch).

## Abstract

The two Condensin complexes in human cells are essential for mitotic chromosome structure. We used homozygous genome editing to fluorescently tag Condensin I and II subunits and mapped their absolute abundance, spacing and dynamic localization during mitosis by fluorescence correlation spectroscopy-calibrated live cell imaging and super-resolution microscopy. While ∼35,000 Condensin II complexes are stably bound to chromosomes throughout mitosis, ∼195,000 Condensin I complexes dynamically bind in two steps, in prometaphase and early anaphase. The two Condensins rarely co-localize at the chromatid axis, where Condensin II is centrally confined but Condensin I reaches ∼50% of the chromatid diameter from its center. Based on our comprehensive quantitative data, we propose a three-step hierarchical loop model of mitotic chromosome compaction: Condensin II initially fixes loops of a maximum size of ∼450 kb at the chromatid axis whose size is then reduced by Condensin I binding to ∼90 kb in prometaphase and ∼70 kb in anaphase, achieving maximum chromosome compaction upon sister chromatid segregation.

## Introduction

A fundamental structural and functional change of the human genome is the compaction of replicated interphase chromatin into rod-shaped mitotic chromosomes. This process of mitotic chromosome condensation is essential for faithful genome partitioning (Hudson et al., 2009) and involves two conserved structural maintenance of chromosomes (SMC) protein complexes, Condensin I and II (Hirano and Mitchison, 1994; Hirano et al., 1997; Ono et al., 2003; Strunnikov et al., 1995; Yeong et al., 2003). Condensins consist of two shared subunits (SMC2 and SMC4) and three isoform-specific subunits - a kleisin (CAP-H or CAP-H2) and two HEAT-repeat proteins (CAP-D2 or CAP-D3 and CAP-G or CAP- G2). SMC2 and SMC4 are back-folded into long coiled-coils, bringing their N and C termini together into two ATPase domains, and are connected at their central domains, creating a “hinge” between the two subunits. The ATPase domains are bridged by the kleisin and associated HEAT-repeat subunits to form a pentameric ring-like architecture with an estimated length of overall ∼60 nm for the human complexes (Anderson et al., 2002). The kleisin and HEAT-repeat subunits have recently been shown to bind DNA in a unique safety belt arrangement (Kschonsak et al., 2017) and the complexes can progressively move on DNA as motors *in vitro* (Terakawa et al., 2017), which is consistent with the hypothesis that they actively form and stabilize DNA loops (Alipour and Marko, 2012; Goloborodko et al., 2016A,B; Nasmyth, 2001).

Within the cell, Condensin II is located in the nucleus and has access to chromosomes throughout the cell cycle, whereas Condensin I is cytoplasmic during interphase and can only localize to mitotic chromosomes after nuclear envelope breakdown (NEBD) in prometaphase (Gerlich et al., 2006; Hirota et al., 2004; Ono et al., 2003; Ono et al., 2004). Consistent with this distinct sub-cellular localization, RNAi and protein depletion experiments have proposed that the two Condensin isoforms promote different aspects of mitotic chromosome compaction, with Condensin II promoting axial shortening in prophase and Condensin I compacting chromosomes laterally in prometa- and metaphase (Green et al., 2012; Hirota et al., 2004; Ono et al., 2003; Ono et al., 2004). Both Condensins localize to the longitudinal axis of mitotic chromosomes and are part of the insoluble non-histone “scaffold” (Maeshima and Laemmli, 2003; Ono et al., 2003).

Extensive structural, biochemical, cell biological and molecular biological research over the last two decades led to numerous models about how Condensins may shape mitotic chromosomes (reviewed in: Cuylen and Haering, 2011; Hirano, 2012; Hirano, 2016; Kalitsis et al., 2017; Kinoshita and Hirano, 2017; Kschonsak and Haering, 2015; Piskadlo and Oliveira, 2016; Uhlmann, 2016). Condensins have been proposed to make topological linkages between two regions within the same chromatid (Cuylen et al., 2011) and thereby introduce loops in the DNA molecule, which, according to the loop-extrusion theory (Alipour and Marko, 2012; Goloborodko et al., 2016A,B; Nasmyth, 2001) and very recent evidence *in vitro* (Ganji et al., 2018), compact mitotic chromosomes and contribute to their mechanical stabilization (Gerlich et al., 2006; Houlard et al., 2015). However, how such Condensin-mediated linkages could organize the hundreds of megabase-sized DNA molecule of a human chromosome and how Condensin I and II mediate different aspects of the overall compaction process is still poorly understood. A key requirement to formulate realistic mechanistic models is to know the copy number and stoichiometry as well as the precise spatial arrangement of Condensin I and II within a mitotic chromatid. However, such quantitative data about Condensins in single dividing cells are currently missing.

To address this gap in our knowledge, we set out to quantitatively determine the dynamic association of Condensin I and II with chromosomes throughout mitosis and resolve their spatial organization relative to the axis of single chromatids. To this end, we took advantage of genome editing in human cells to create homozygous fluorescent knock-ins for SMC, kleisin and HEAT-repeat subunits of both Condensins. We then employed fluorescence correlation spectroscopy (FCS)-calibrated live cell imaging to determine the number of Condensin subunits on chromosomes over the course of mitosis, as well as stimulated emission depletion (STED) super-resolution microscopy of single chromatids in specific mitotic stages to investigate the axial organization of Condensin complexes. By measuring the physical and normalizing for the genomic length of mitotic chromosomes, our comprehensive quantitative imaging-based data of human Condensin complexes on mitotic chromosomes allow for the first time to formulate models of mitotic chromosome structure which are based on concentrations, physical distances and average genomic spacing of the key protein complexes that structure DNA during mitosis. Based on these data, we propose a three-step hierarchical loop formation model for mitotic chromosome compaction that makes quantitative predictions about the hierarchy and size of loops formed in the chromosomal DNA to achieve accurate genome partitioning during cell division.

## Results and discussion

### Quantitative imaging of five Condensin subunits

To quantitatively investigate the dynamic subcellular distribution of Condensin complexes during mitosis, we generated genome-edited human HeLa Kyoto cell lines in which all alleles of the endogenous genes for the shared subunit SMC4 and two pairs of kleisin and HEAT-repeat subunits specific to Condensin I (CAP-H and CAP-D2) or Condensin II (CAP-H2 and CAP-D3) were tagged with mEGFP (Fig. S1A,B). Specific and homozygous tagging was verified with a multi-step quality control pipeline (Koch et al., 2017 *Preprint*; Mahen et al., 2014; for details see Materials and Methods), ensuring that the tagged subunit was expressed at physiological levels, incorporated into complexes during interphase and mitosis (Fig. S1B) and that mitotic progression and timing were not perturbed (Fig. S1C), thus indicating full functionality of the fusion proteins.

We then applied automated FCS-calibrated confocal time-lapse microscopy to image Condensin subunits during mitosis relative to spatio-temporal landmarks that defined cell (extracellular Dextran-Dy481XL) and chromatin volumes (DNA stained by SiR-DNA; Cai et al., 2017 *Preprint*; Lukinavičius et al., 2015). This calibrated 4D imaging data allowed us to derive time-resolved 3D maps of subcellular protein concentration and copy number of Condensin subunits (Fig. 1A; Fig. S1D-I; Politi et al., 2017 *Preprint*) and distinguish the soluble cytoplasmic from the chromatin-bound protein fractions for all mitotic stages until nuclear envelope reformation (Fig. 1B; Cai et al., 2017 *Preprint*; for details see Materials and Methods). Using computational temporal alignment (Cai et al., 2017 *Preprint*), we integrated the data for all Condensin subunits, allowing us to perform a quantitative comparison of their association with mitotic chromosomes (Fig. 1C).

**Fig. 1:**
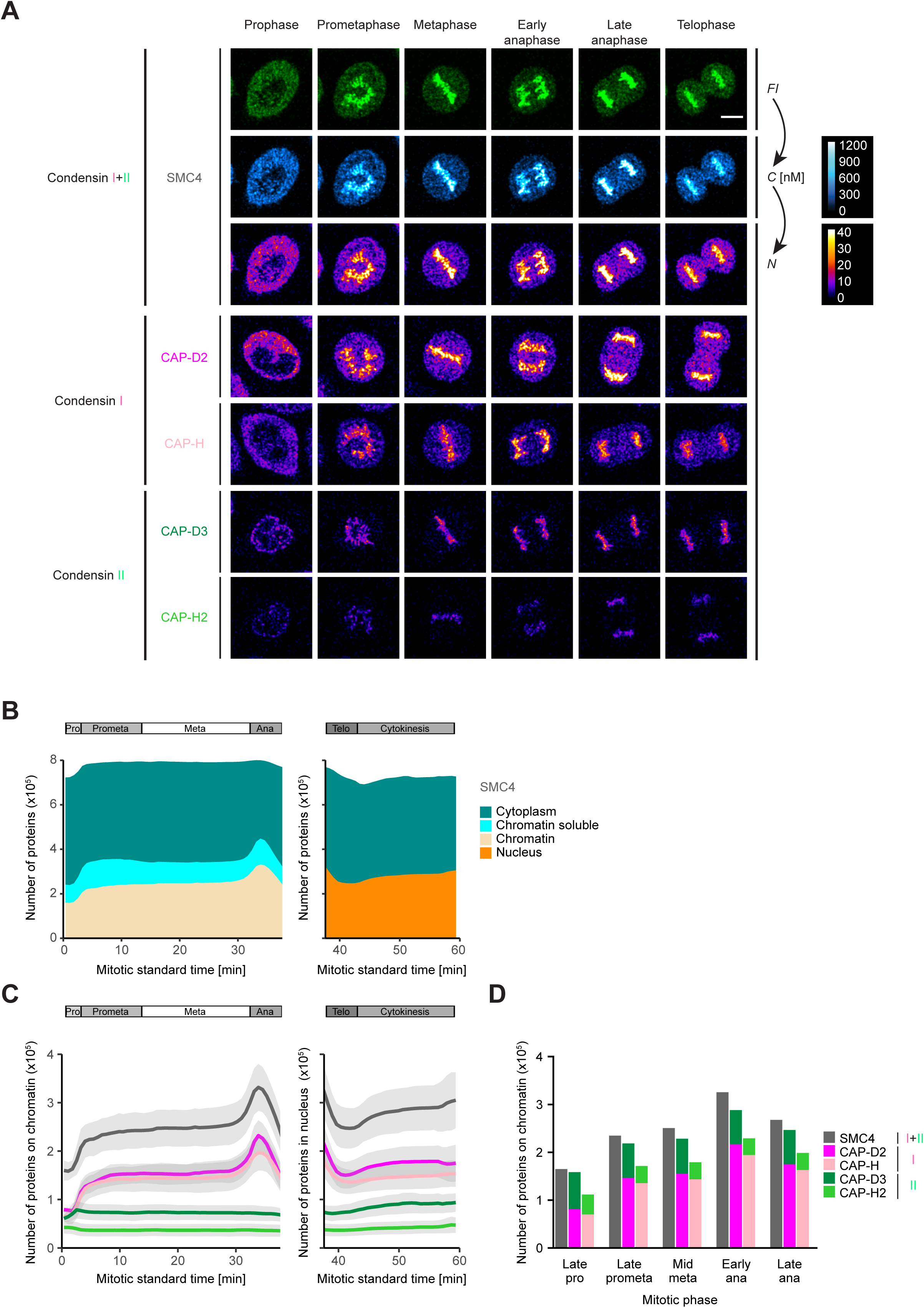
Condensin I and II display different quantitative dynamics during mitosis as determined by automated FCS-calibrated confocal 3D time-lapse imaging. **(A)** Genome-edited HeLa Kyoto cells with homozygously mEGFP-tagged Condensin subunits and chromosomes stained by SiR-DNA were imaged every 90 s for a total of 60 min by 3D confocal microscopy, automatically triggered after prophase onset. Images were calibrated by FCS to convert fluorescence intensities (*FI*) into cellular protein concentration maps (*C*) and cellular protein number maps (*N*) (for details see Fig. S1 D-I). For SMC4 (Condensin I/II) fluorescence intensities, cellular protein concentration and protein number maps are shown. For CAP-D2 and CAP-H (Condensin I) and CAP-D3 and CAP-H2 (Condensin II) protein number maps are depicted. Single *z*-planes are shown for specific mitotic phases comparable between subunits. Condensin II is nuclear/on chromosomes throughout mitosis, Condensin I is cytoplasmic in prophase and gains access to mitotic chromosomes after NEBD in prometaphase. A Gaussian blur (σ = 1) was applied to the images for presentation purposes. Scale bar: 10 µm. **(B)** SMC4 protein numbers in specific compartments of mitotic HeLa cells (cytoplasm (turquoise), nucleus (orange), chromatin-bound (beige), chromatin soluble (blue)) are plotted against the mitotic standard time and the corresponding mitotic phases are indicated. Shown is the average of 17 cells from 4 independent experiments. Cellular landmarks were used for 3D segmentation and conversion of compartment-specific protein concentrations into protein numbers (for details see Fig. S1I and Materials and Methods). Chromosome-bound and soluble chromatin proteins defined as the freely diffusing proteins within the chromatin volume were distinguished until anaphase as described in Materials and Methods. For late mitotic stages the nuclear compartment could not be divided into chromatin-bound and soluble proteins. **(C)** The numbers of chromosome-bound (pro- to anaphase) and nuclear Condensin subunits (telophase and cytokinesis) are plotted against the mitotic standard time: Condensin II subunits (CAP-D3, dark green; CAP-H2, light green), Condensin I subunits (CAP-D2, dark magenta; CAP-H, light magenta) and shared SMC4 (dark grey). Mean (colored line) and SD (light grey area) are shown. **(D)** The number of chromosome-bound Condensin subunits from (C) are plotted for selected mitotic stages according to the images shown in (A). SMC4 subunits (grey) as well as the sum of corresponding HEAT-repeat subunits (CAP-D2, dark magenta, Condensin I; CAP-D3, dark green, Condensin II) and corresponding kleisin subunits (CAP-H, light magenta, Condensin I; CAP-H2, light green, Condensin II) are shown. For (C) and (D) the mean of ∼20 cells per subunit (range: 10-36) from 3-7 independent experiments is plotted; 36, 22, 10, 17 and 17 cells from 7, 3, 5, 3, and 4 experiments for CAP-H, CAP- H2, CAP-D2, CAP-D3 and SMC4, respectively.

### About 200,000 Condensin I complexes bind in two steps to mitotic chromosomes

We first examined the total number of all Condensins (SMC4) as well as the number of Condensin I (CAP-D2/CAP-H) and Condensin II (CAP-D3/CAP-H2) subunits in the whole cell (Fig. S2A), in the cytoplasm (Fig. S2B) and on chromosomes (Fig. 1C) throughout mitosis. In the whole cell, the number of Condensin I (∼675,000 for CAP-D2) and Condensin II (∼115,000 for CAP-D3) and correspondingly also Condensin I plus II molecules (∼780,000 for SMC4; Fig. 1B, Fig. S2A) remained constant throughout mitosis, which indicates that neither new protein synthesis nor protein degradation are used to regulate their function during cell division.

However, the number of Condensin I subunits associated with mitotic chromosomes changed dramatically (Fig. 1C). With the major fraction gaining access to chromosomes only upon NEBD, the level of chromosome-bound Condensin I reaches a first plateau during prometa- and metaphase, with ∼150,000 CAP-D2 molecules per replicated genome. Upon anaphase onset, a second wave of Condensin I subunits binds mitotic chromosomes, which results in a maximum value of ∼215,000 CAP-D2 molecules on all chromosomes. After chromosome segregation has been accomplished, the number of Condensin I subunits declines rapidly to ∼125,000 CAP-D2 molecules at telophase. Condensin II subunits, by contrast, are much less abundant on chromosomes, with only ∼70,000 CAP-D3 molecules per replicated genome, a number that stays constant throughout mitosis (Fig. 1C).

Although we recorded each subunit in individual homozygous knock-in cell lines, the numbers of chromatin-localized isoform-specific subunits (e.g. ∼150,000 CAP-D2 (Condensin I) and ∼70,000 CAP-D3 (Condensin II) in metaphase) summed up very close to the number of chromosome-bound copies of the shared SMC4 subunit (∼250,000 in metaphase) throughout mitosis (Fig. 1D), validating our cell line generation and quantitative imaging pipelines. The number of ∼250,000 SMC4 subunits on metaphase chromosomes corresponds to a concentration of ∼520 nM on chromatin (Fig. S2C), while the cytoplasmic concentration of ∼190 nM from the ∼530,000 SMC4 molecules in this larger compartment is lower than the chromosomal concentration (Fig. S2D).

As Condensin I is more abundant than Condensin II, the two-step binding observed for its kleisin and HEAT-repeat subunits is also reflected in the dynamics of the shared SMC4 subunit (Fig. 1B,C). This behavior is in line with previous reports based on immunostaining or overexpression (Gerlich et al., 2006; Hirota et al., 2004; Ono et al., 2003, Ono et al., 2004).

Our quantitative dynamic imaging data suggests that Condensin II is bound to mitotic chromosomes in prophase and might promote the initial mitotic chromosome compaction, while Condensin I is mostly cytoplasmic (Fig. 1A; Gerlich et al., 2006). The majority of Condensin I rapidly localizes to chromosomes in prometaphase and in a second step in anaphase (Gerlich et al., 2006; Mora-Bermúdez et al., 2007), when it likely promotes further compaction of mitotic chromatids. In the future it will be very interesting to study how the two waves of Condensin I binding to mitotic chromosomes from a constant cellular pool are regulated during mitosis.

### The kleisin subunit appears to be limiting for Condensin holocomplexes on chromosomes

Interestingly, while showing very similar dynamics, the number of kleisin subunits for Condensin I and II (CAP-H and CAP-H2) cell lines was consistently lower than the number of the corresponding HEAT-repeat subunits (CAP-D2 and CAP-D3; Fig. 1C, D), indicating that some chromosome-localizing SMC and HEAT-repeat subunits are missing kleisin subunits to close the pentameric ring, as also suggested in earlier biochemical studies (Kimura et al., 2001; Ono et al., 2003). We confirmed this with both C- and N-terminally tagged CAP-H/H2 subunits and several homozygous clones (Fig. S2F,G; Cai et al., 2017 *Preprint*). For Condensin II, about half of the chromosome-bound CAP-D3 subunits could form holocomplexes (Fig. S2G), while for Condensin I ∼90% of the CAP-D2 subunits would find sufficient kleisin binding partners for holocomplexes (Fig. S2F). Based on the limiting kleisin subunits, we estimate a maximal number of ∼35,000 Condensin II and ∼195,000 Condensin I holocomplexes on anaphase chromosomes (Fig. 1C) and will use the kleisin subunits as indicators for Condensin holocomplexes in the following. Our finding that there are about twice as many chromosome-localizing CAP-D3 subunits than CAP-H2 subunits suggests that either the stoichiometry of CAP-D3 to CAP-H2 in Condensin II complexes is higher than two to one or that CAP-D3 can bind mitotic chromosomes independently of being part of Condensin II complexes. Consistent with the latter possibility, precipitating CAP-H2-mEGFP from our endogenously tagged cell lines revealed an unbound fraction of CAP-D3 in the supernatant.

### Condensin I is up to six times more abundant on mitotic chromosomes than Condensin II

Comparison of the number of kleisin subunits as indicators for the maximum amount of holocomplexes at the same mitotic stage revealed that the ratio of Condensin I to Condensin II is about 4:1 on metaphase chromosomes and increases to about 5.6:1 in early anaphase (Fig. 1C), which is consistent with an earlier biochemical assessment in *Xenopus* egg extracts (Ono et al., 2003). The much higher abundance and mitosis-specific binding during the two major stages of compaction suggest that Condensin I is the major contributor to the formation of mitotic chromosomes, consistent with it being required and, together with topoisomerase II, sufficient for mitotic chromosome assembly *in vitro* (Shintomi et al., 2015; Shintomi et al., 2017).

Overall, our quantitative FCS-calibrated live cell imaging data based on homozygous endogenously tagged Condensin subunits revealed that Condensin I is 1.6-5.6 times more abundant than Condensin II during mitosis and binds to mitotic chromosomes in two steps in prometaphase and early anaphase. In contrast, the less abundant Condensin II does not change its association with chromosomes during mitosis, suggesting very different roles for the two Condensin complexes in the structural organization of mitotic chromosomes.

### Condensin I interacts dynamically with mitotic chromosomes, while Condensin II binds stably

To investigate how stable the two Condensins are bound to mitotic chromosomes, we investigated their residence times by FRAP (Fig. S2H-J; for details see Materials and Methods). This revealed that the step-wise association of Condensin I in prometaphase and anaphase and its subsequent rapid decrease in telophase is underpinned by a relatively short chromosomal residence time of ∼2 min (Fig. S2K) and a low immobile fraction (<20%; Fig. S2L) on metaphase chromosomes. By contrast, Condensin II is more stably bound, with a longer residence time (>5 min; Fig. S2K) and a much higher immobile fraction (>60%; Fig. S2L), consistent with its constant abundance on mitotic chromosomes (Fig. 1C,D). Within 10 min of metaphase ∼85% of chromosome-bound Condensin I and ∼30% of chromosome-bound Condensin II are exchanged. These findings with endogenously tagged Condensins confirm previous reports based on overexpressed proteins (Gerlich et al., 2006). The high turnover of Condensin I binding on mitotic chromosomes is in line with a relatively high cytoplasmic fraction (>60% of the total cellular protein; Fig. S2E) in contrast to Condensin II, for which the large majority is bound to chromosomes (Fig. S2E) and exchanges much more slowly. Interestingly, this behavior is consistent between CAP-H2 and CAP-D3. Since the abundance measurements and immunoprecipitations indicate that some CAP-D3 proteins can bind chromosomes without being associated with Condensin II complexes (Fig. 1C,D; Fig. S2F), this fraction would then bind with a residence time that cannot be distinguished from CAP-H2 by FRAP. The slow binding dynamics and stable abundance of Condensin II on mitotic chromosomes (Fig. 1C,D) suggest a more structural and stabilizing role, while the dynamic step-wise binding and dissociation of Condensin I is consistent with it playing an actively regulated role in both mitotic compaction and de-compaction of chromosomes.

### Super-resolution imaging of Condensin I and II in prometaphase and anaphase

Since Condensin I and II display very different chromosome binding dynamics and stoichiometries in mitosis, we investigated whether they also differ in their sub-chromosomal localization. To this end, we used super-resolution (STED) microscopy to acquire 3D images and localize our tagged Condensin subunits relative to DNA within single chromatids in cells fixed in specific mitotic stages (Fig. S3A; for details see Materials and Methods). We focused on late prometaphase because at this stage Condensin I has reached its first binding plateau and sister chromatids are already resolved (Nagasaka et al., 2016; Fig. 1C), as well as on early anaphase when Condensin I abundance peaks (Fig. 1C) and chromatids reach their maximum compaction (Mora-Bermúdez et al., 2007). Using exactly the same labeling and imaging conditions to visualize the different Condensin subunits allowed us to quantitatively compare and integrate the data between the two isoforms.

Our STED data revealed that both Condensins localized along the longitudinal axis in the center of chromatid arms in both prometa- and anaphase (Fig. 2), consistent with earlier reports (Maeshima and Laemmli, 2003; Ono et al., 2003; Poonperm et al., 2015). In prometaphase, Condensin I decorated chromatids more densely than Condensin II and became even more densely localized in anaphase, consistent with Condensin I’s higher abundance and increased binding as observed by FCS-calibrated imaging in live cells (Fig. 1C,D).

**Fig. 2:**
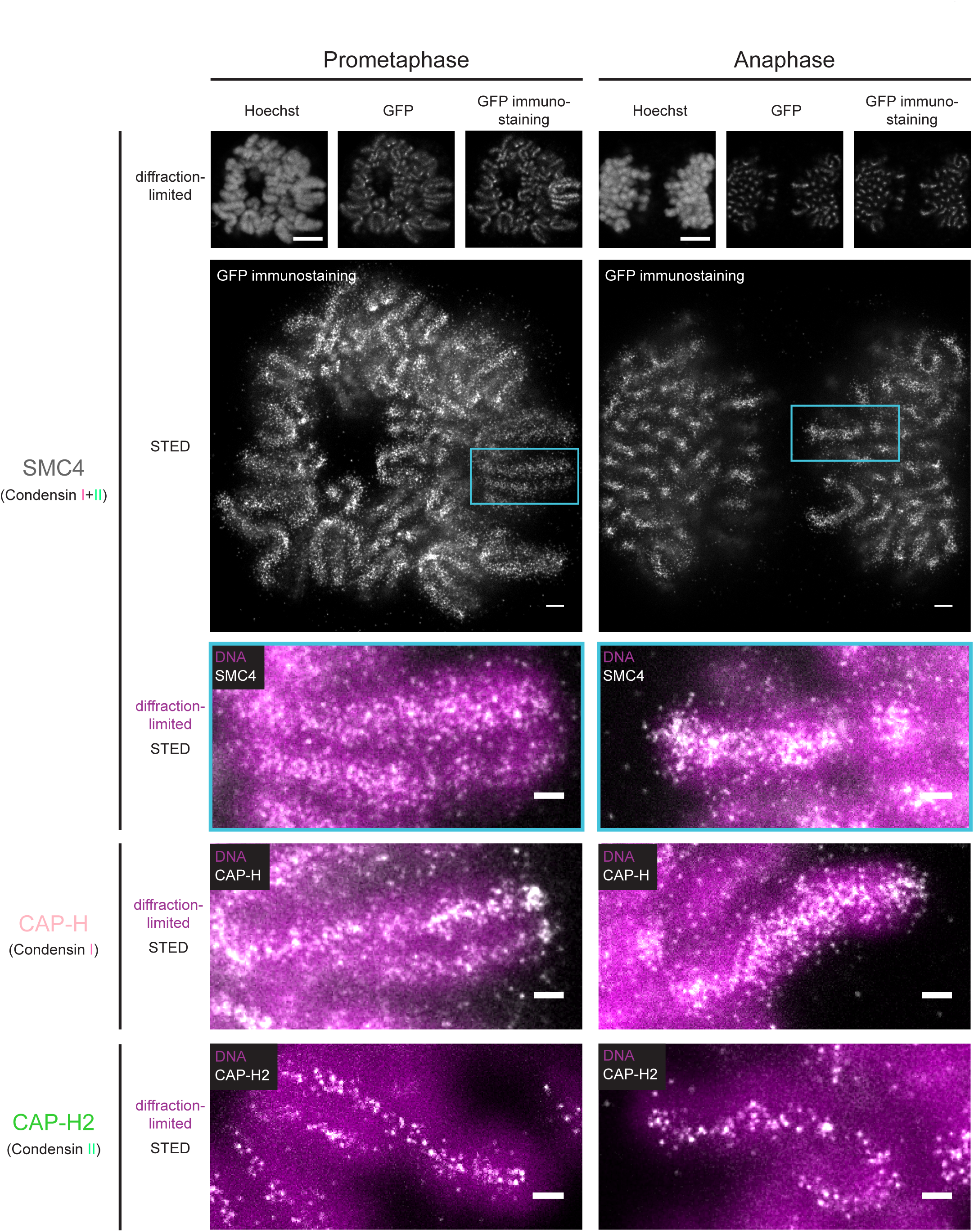
STED super-resolution imaging of Condensin I and II. Condensin subunits were immunostained (anti-GFP) and mitotic cells with chromatids oriented in parallel to the focal plane were selected for imaging. DNA (Hoechst) and Condensins (mEGFP tag and anti-GFP immunostaining) were imaged diffraction-limited, while immunostained Condensin was also imaged by STED microscopy. Representative images of mitotic chromatids with SMC4 (Condensin I + II), CAP-H (Condensin I) and CAP-H2 (Condensin II) in late prometaphase (left) and early anaphase (right) are shown. For SMC4 (upper panels) whole-cell overview images as well as overlays of super-resolved Condensin (white) and diffraction-limited DNA (magenta) imaging are shown. Overlays represent zoom-ins into single chromatids (turquoise box). For CAP-H (middle panels) and CAP-H2 (lower panels) only overlays are depicted. Representative images of single *z* planes are shown. Scale bars: 5 µm (whole cell diffraction-limited), 1 µm (whole cell STED), 500 nm (zoom-ins).

### Condensin II is more confined to the chromatid axis than Condensin I

To quantify how far away from the chromatid center Condensins are localized, we used computational image analysis to determine the average full width at half maximum (FWHM) of Condensin subunits relative to DNA (Fig. S3A,B; for details see Materials and Methods). Condensin II is significantly more restricted to the center of the chromatid, occupying ∼30-35% of the ∼1,100 nm diameter of the chromatid arm (Fig. 3A). By contrast, Condensin I localization can be detected at up to 50% of the chromatid diameter in both prometa- and anaphase (Fig. 3A). Interestingly, three out of four isoform-specific subunits display a statistically significant increase (∼100 nm) in their FWHM from prometaphase to anaphase, consistent with the idea that the final chromatid arm compaction (Mora-Bermúdez et al., 2007) results in a widening of the internal chromosome scaffold.

**Fig. 3:**
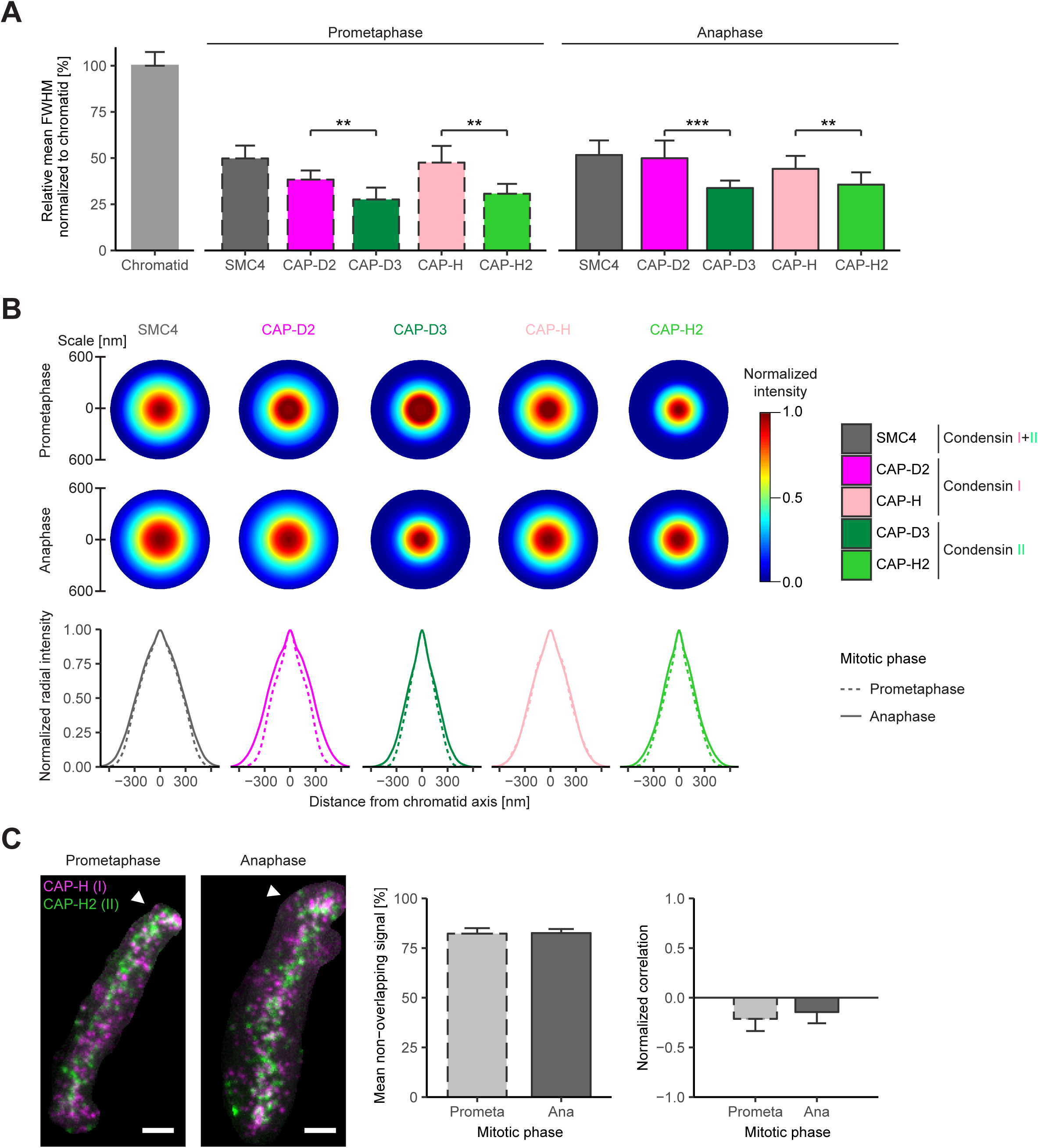
Non-overlapping localization of Condensin I and II along the mitotic chromatid axis. **(A)** The full width at half maximum (FWHM) of Condensin subunit distribution and chromatid arms was analyzed (for details see Fig. S3 A,B). Condensin subunit FWHM are plotted relative to the chromatid FWHM in two mitotic phases (mean + SD; ∼13 chromatids; *n* = 5-20; prometaphase: *n* = 7, 12, 5, 13 and 20 chromatids for CAP-H, CAP-H2, CAP-D2, CAP-D3 and SMC4, respectively; anaphase: *n* = 15, 16, 11, 20 and 15 chromatids for CAP-H, CAP-H2, CAP-D2, CAP-D3 and SMC4, respectively): prometa-(dashed line, left panel) and anaphase (solid line, right panel). Student’s two-tailed t-test: in prometaphase *p* = 0.0034 for CAP-D2/CAP-D3 and *p* = 0.0014 for CAP- H/CAP-H2; in anaphase *p* = 0.0002 for CAP-D2/CAP-D3 and *p* = 0.0016 for CAP- H/CAP-H2. Comparison of FWHM for the same Condensin subunit between prometa- and anaphase: CAP-D2 (*p* = 0.0067), CAP-D3 (*p* = 0.0064), CAP-H (*p* = 0.3966), CAP- H2 (*p* = 0.0391). * *p*<0.05, ** *p*<0.01, *** *p*<0.001. Since the chromatid FWHM between prometaphase and anaphase was not significantly different, all chromatids were pooled for all subunits and mitotic phases (∼1.1 µm; *n* = 134). **(B)** Radial intensity distribution of Condensin subunits on chromatid arms (for details see Fig. S3C). *xy* Condensin intensity profiles were symmetrized, reconstructed as radially symmetric distribution and centrally aligned with the chromatid. Condensin intensities were peak-normalized per subunit and mitotic phase and are displayed within circles representing the chromatid width determined in (A) as diameter. Circular radial intensity profiles are shown for all analyzed Condensin subunits in prometaphase and anaphase (upper panel). The *z*-projected (sum) Condensin intensity profiles are plotted for prometa-(dashed line) and anaphase (solid line) together for each subunit (lower panel). Averaged intensity profiles among all chromatids per subunit and mitotic stage are plotted (∼13 chromatids; *n* = 5-20). **(C)** Immunostaining of double homomozygously Condensin-tagged cell lines: Condensin I (CAP-H-Halo; magenta) and II (CAP-H2- mEGFP; green). One *z* plane of a representative prometaphase and anaphase chromatid is shown and telomeres are indicated by white arrowheads (left panel). Scale bars: 500 nm. While the labeling efficiencies of the two tags make a quantitative comparison of the protein abundances difficult, they were sufficiently high to allow us to test whether the two isoforms co-localize at the same chromosomal loci. Thus, Condensin I and II signals were segmented and analyzed for co-localization. Mean percentage of non-overlapping segmented voxels between Condensin I and II is plotted (82% in both prometa- and anaphase; middle panel). Normalized correlation of Condensin I and II signals is plotted (anti-correlated in both prometa- and anaphase; right panel). Mean + SD (prometaphase: *n* = 19; anaphase: *n* = 18).

To determine where most of the Condensin molecules are located within the chromatid width, we quantified their radial intensity distributions perpendicular to the longitudinal chromatid axis (Fig. S3C; for details see Materials and Methods). Consistent with the FWHM measurements, Condensin I subunits display a broader intensity distribution than Condensin II both in prometa- and anaphase (Fig. 3B). 90% of the Condensin I (CAP-D2) signal is contained within a diameter of 560 nm in prometaphase, extending to 760 nm in anaphase, while 90% of Condensin II (CAP-D3) were confined closer to the chromatid axis (480 nm and 560 nm diameter in prometaphase and anaphase, respectively). Interestingly, the shape of the radial intensity profile also differs between the isoforms. Condensin II subunits show a narrow peak at the chromatid axis, consistent with an enrichment in the chromatid center, whereas Condensin I subunits show a more bell-shaped profile with wider shoulders, consistent with the conclusion that many Condensin I complexes localize further away from the central chromatid axis (Fig. 3B). This bell-like distribution of Condensin I broadens from prometaphase to anaphase (Fig. 3B), indicating that most of the Condensin I mass that is added in anaphase binds distally from the central chromatid axis.

### Condensin I and II are not co-localized on mitotic chromatid arms

The differences in average axial distribution of Condensin I and II suggest that the two complexes at least partially act at different loci on the chromosomal DNA molecule. To test this hypothesis directly, we imaged Condensin I and II in the same cell in 3D by two-color STED microscopy, which has a lateral resolution on the same scale as expected for the length of single Condensin complexes (Anderson et al., 2002). To this end, we generated a double homozygous knock-in cell line in which we tagged CAP-H and CAP- H2 differentially with a Halo and a mEGFP tag. Remarkably, co-imaging both kleisin subunits showed that Condensin I and II signals were anti-correlated, with ∼82% of the spot-like signals showing no spatial overlap in prometaphase or in anaphase (Fig. 3C; for details see Materials and Methods). The few co-localization events that we detected were enriched at centromeres and telomeres, where Condensins reach very high concentrations (Fig. 3C, arrowheads), consistent with previous reports based on diffraction-limited imaging (Gerlich et al., 2006; Oliveira et al., 2005; Ono et al., 2004; Ribeiro et al., 2009). Interestingly, our super-resolved images indicate an alternating localization of Condensin I and II along the chromatid arm, in line with previous studies which could not resolve single complexes (Maeshima and Laemmli, 2003; Ono et al., 2003). However, our higher resolution data do not show quantifiable regular or periodic patterns.

In summary, our super-resolution analysis of mitotic chromosomes in whole human cells shows that Condensin I and II differ in their localization within the chromatid, with Condensin II being confined to the axis while Condensin I also binds more peripherally, as also shown for *in vitro* reconstituted chromatids (Shintomi et al., 2017), and that they do not bind to the same sites on mitotic chromatid arms.

### Genomic and physical spacing of Condensin subunits on mitotic chromosomes

In order to use our quantitative data of Condensin abundance, binding and sub-chromosomal position to formulate a model for how chromosomal DNA molecules might be structured and compacted in mitosis, we needed to establish the relationship between physical distances and genomic lengths for mitotic chromosomes. While the average genome size of the HeLa Kyoto genome is 7.9 billion bp (on average hypotriploid, e.g. ∼64 chromosomes; Landry et al., 2013), its physical dimensions have not yet been determined. We therefore measured the total lengths of all chromatid axes per HeLa cell in different mitotic stages by tracing the SMC4-mEGFP labeled chromatid axis relative to DNA in high resolution 3D data sets collected from live cells (Fig. 4A; for details see Materials and Methods). This approach avoids any fixation artefacts and thus enables direct comparison to our live cell abundance measurements. Consistent with an increase in chromatid compaction during mitosis (Hériché et al., 2014, Mora-Bermúdez et al., 2007), the total chromatid length decreased from ∼1,300 µm in prometaphase to ∼1,150 µm in metaphase, reaching ∼925 µm in the most compacted state in anaphase (Fig. 4B). Thus, an “average” HeLa chromatid of 123 Mb has a physical length of ∼9 µm in metaphase and contains ∼1,100 Condensin I (CAP-H) and ∼270 Condensin II (CAP-H2) holocomplexes (Table 1). During sister chromatid segregation in anaphase, the number of Condensin I holocomplexes increases to ∼1,500 and the chromatid is shortened to ∼7 µm (Table 1).

**Fig. 4:**
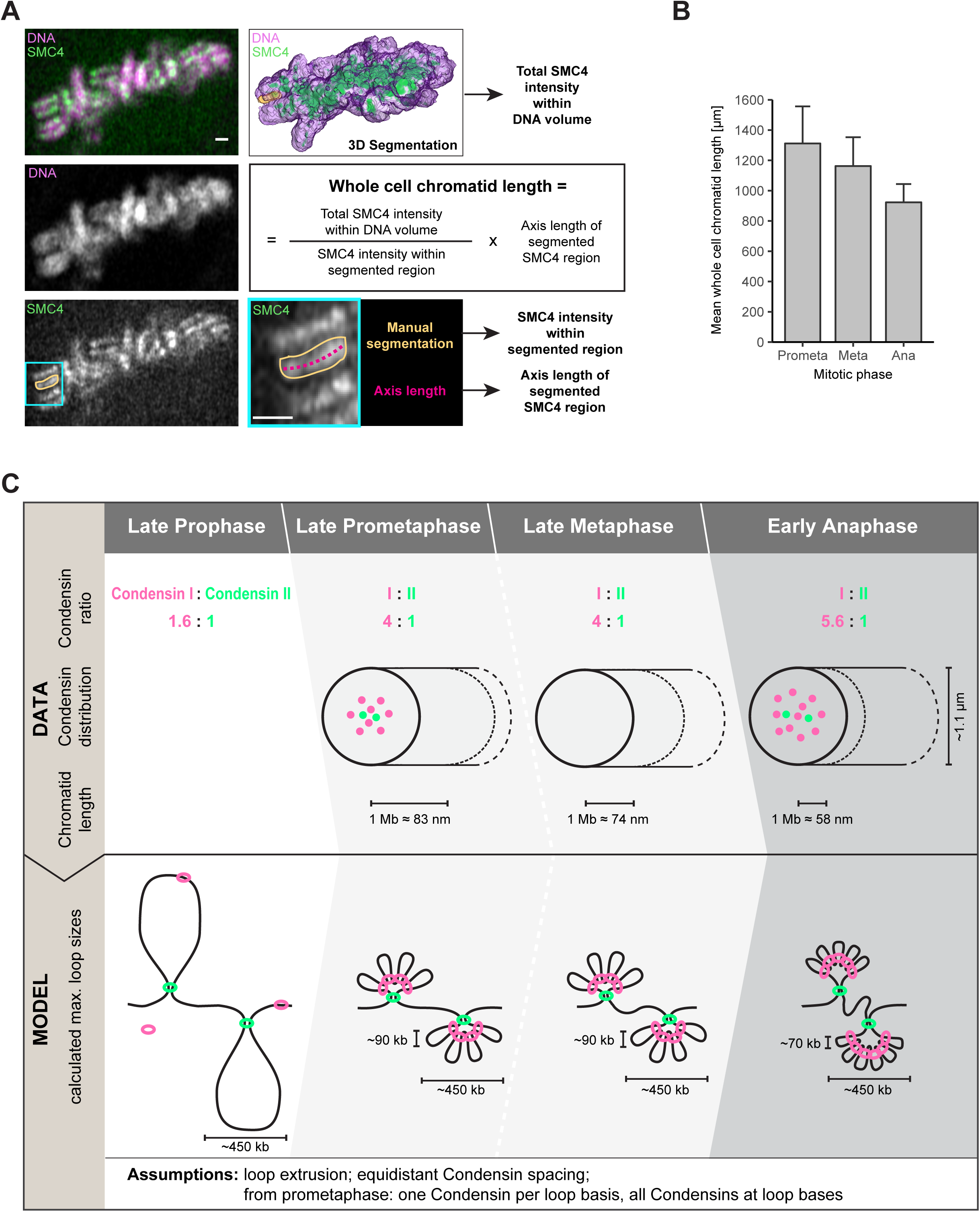
A three-step hierarchical looping model of mitotic chromosome compaction based on a comprehensive 3D map of Condensins in human mitotic cells. **(A,B)** Whole chromatid length in mitotic HeLa Kyoto cells. **(A)** High-resolution *z*-stacks of SMC4-tagged live cells stained with SiR-DNA to visualize chromosomes were acquired. For each cell, the total Condensin (green) and chromosome (magenta) volume was segmented in 3D and the total Condensin intensity within the chromosome volume was determined. In addition, for each cell several (3-5) clearly recognizable stretches of Condensin signal along the axis of chromatid arms were segmented manually and their Condensin intensity and corresponding axis length determined. These were used to compute the mean chromatid length per cell according to the formula indicated (see also Materials and Methods). Scale bar: 1 µm. **(B)** Whole cell chromatid length for late prometaphase, metaphase and early anaphase (mean + SD; *n* = 3 cells). **(C)** The quantitative 3D map of Condensins in human mitotic HeLa Kyoto cells suggests a Condensin-based three-step hierarchical looping model of mitotic chromosome compaction. Step 1: Condensin II, which is present in the nucleus in interphase, promotes axial compaction in prophase. This compaction results in maximum ∼450 kb long loops if fully extruded, with Condensin II localizing within the central 30-35% of the mitotic chromatid diameter (∼1.1 µm). Step 2: After NEBD in prometaphase, the majority of the in interphase and prophase mainly cytoplasmic Condensin I gains access to mitotic chromosomes and binds to the central chromatid axis but also reaches more peripheral sites up to 50% of the chromatid diameter. The first phase of Condensin I binding leads to a 4-fold higher abundance compared to Condensin II on metaphase chromosomes and drives mainly lateral compaction in prometaphase and metaphase by subdividing the large Condensin II loops 4 times into sub-loops of a size of ∼90 kb. In this step, the chromatid axis length constraining 1 Mb DNA decreases to 83 nm in prometa- and further to 74 nm in metaphase. This Condensin I-driven sub-loop formation could in principle start already before NEBD based on the minor Condensin I fraction present on prophase chromosomes. Step 3: Upon anaphase onset, a second wave of Condensin I binds to mitotic chromosomes, reaching a 5.6-fold higher abundance than Condensin II and localizing even further away from the chromatid axis. These additional Condensin I complexes drive the final axial shortening step by further subdividing Condensin II loops into 5-6 sub-loops, leading to a final loop size of ∼70 kb and 1 Mb DNA being now constraint to 58 nm chromatid axis length when maximal chromosome compaction is reached at the end of anaphase. The Condensin abundances and spatial dimensions in the upper half of the model were determined in this work (Condensin I:II ratio; chromatid width occupied by Condensin I/II; relation of genomic and physical genome length). The loop sizes in the lower half of the model calculated from these data are based on an equidistant Condensin spacing along the genome and the assumption of fully extruded loops according to the loop-extrusion theory, whereby each loop basis would be occupied by one Condensin holocomplex and all Condensin holocomplexes are at loop bases (for prophase only for Condensin II, from prometaphase to anaphase for both Condensin complexes). However, the DNA path and spatial arrangement of loops in the lower half of the model is drawn hypothetically to represent the determined DNA length.

**Table 1:**
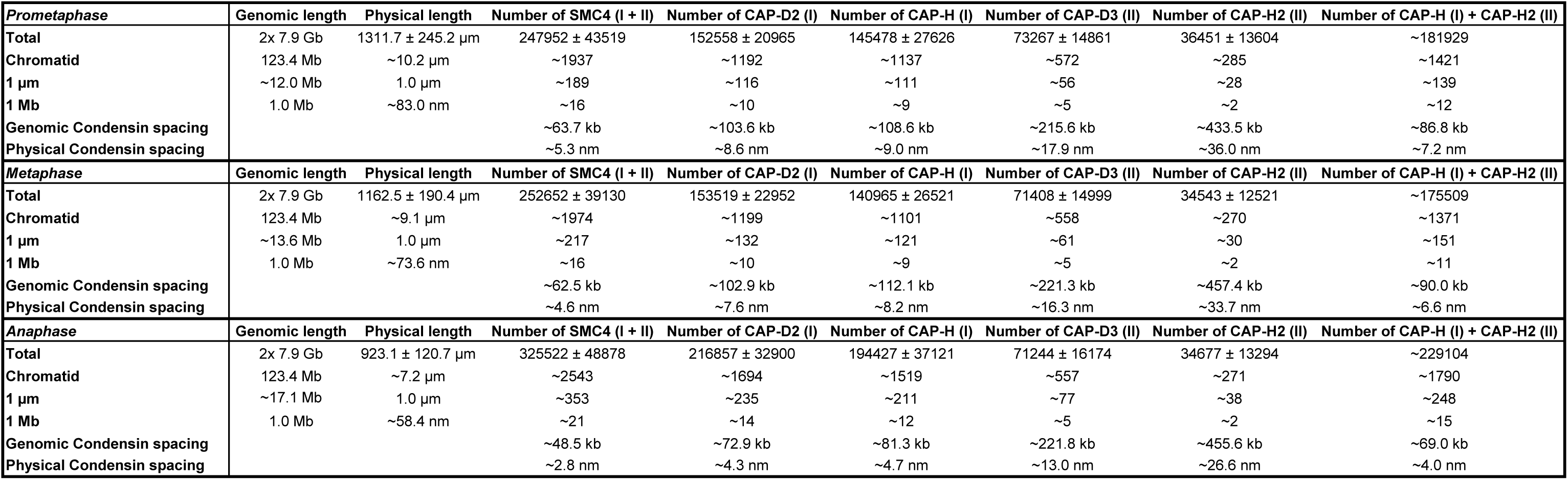
Genomic and physical DNA length of an average HeLa Kyoto cell, the absolute amount of Condensin subunits on mitotic chromosomes and their linear spacing along the chromatid axis in late prometa-, meta- and early anaphase. Calculations are based on the 7.9 billion bp HeLa Kyoto genome with on average 64 chromosomes (Landry et al., 2013), the whole cell chromatid lengths for late prometaphase, metaphase and early anaphase, as well as the numbers of Condensin subunits bound to chromosomes as determined by FCS-calibrated imaging. The genomic and physical Condensin spacing assumes an equidistant distribution of Condensins along the mitotic chromatid axis.

| *Prometaphase* | Genomic length | Physical length | Number of SMC4 (I + II) | Number of CAP-D2 (I) | Number of CAP-H (I) | Number of CAP-D3 (II) | Number of CAP-H2 (II) | Number of CAP-H (I) + CAP-H2 (II) |
| --- | --- | --- | --- | --- | --- | --- | --- | --- |
| **Total** | 2x 7.9 Gb | 1311.7 ± 245.2 µm | 247952 ± 43519 | 152558 ± 20965 | 145478 ± 27626 | 73267 ± 14861 | 36451 ± 13604 | ~181929 |
| **Chromatid** | 123.4 Mb | ~10.2 µm | ~1937 | ~1192 | ~1137 | ~572 | ~285 | ~1421 |
| **1 µm** | ~12.0 Mb | 1.0 µm | ~189 | ~116 | ~111 | ~56 | ~28 | ~139 |
| **1 Mb** | 1.0 Mb | ~83.0 nm | ~16 | ~10 | ~9 | ~5 | ~2 | ~12 |
| **Genomic Condensin spacing** |  |  | ~63.7 kb | ~103.6 kb | ~108.6 kb | ~215.6 kb | ~433.5 kb | ~86.8 kb |
| **Physical Condensin spacing** |  |  | ~5.3 nm | ~8.6 nm | ~9.0 nm | ~17.9 nm | ~36.0 nm | ~7.2 nm |
| *Metaphase* | Genomic length | Physical length | Number of SMC4 (I + II) | Number of CAP-D2 (I) | Number of CAP-H (I) | Number of CAP-D3 (II) | Number of CAP-H2 (II) | Number of CAP-H (I) + CAP-H2 (II) |
| **Total** | 2x 7.9 Gb | 1162.5 ± 190.4 µm | 252652 ± 39130 | 153519 ± 22952 | 140965 ± 26521 | 71408 ± 14999 | 34543 ± 12521 | ~175509 |
| **Chromatid** | 123.4 Mb | ~9.1 µm | ~1974 | ~1199 | ~1101 | ~558 | ~270 | ~1371 |
| **1 µm** | ~13.6 Mb | 1.0 µm | ~217 | ~132 | ~121 | ~61 | ~30 | ~151 |
| **1 Mb** | 1.0 Mb | ~73.6 nm | ~16 | ~10 | ~9 | ~5 | ~2 | ~11 |
| **Genomic Condensin spacing** |  |  | ~62.5 kb | ~102.9 kb | ~112.1 kb | ~221.3 kb | ~457.4 kb | ~90.0 kb |
| **Physical Condensin spacing** |  |  | ~4.6 nm | ~7.6 nm | ~8.2 nm | ~16.3 nm | ~33.7 nm | ~6.6 nm |
| *Anaphase* | Genomic length | Physical length | Number of SMC4 (I + II) | Number of CAP-D2 (I) | Number of CAP-H (I) | Number of CAP-D3 (II) | Number of CAP-H2 (II) | Number of CAP-H (I) + CAP-H2 (II) |
| **Total** | 2x 7.9 Gb | 923.1 ± 120.7 µm | 325522 ± 48878 | 216857 ± 32900 | 194427 ± 37121 | 71244 ± 16174 | 34677 ± 13294 | ~229104 |
| **Chromatid** | 123.4 Mb | ~7.2 µm | ~2543 | ~1694 | ~1519 | ~557 | ~271 | ~1790 |
| **1 µm** | ~17.1 Mb | 1.0 µm | ~353 | ~235 | ~211 | ~77 | ~38 | ~248 |
| **1 Mb** | 1.0 Mb | ~58.4 nm | ~21 | ~14 | ~12 | ~5 | ~2 | ~15 |
| **Genomic Condensin spacing** |  |  | ~48.5 kb | ~72.9 kb | ~81.3 kb | ~221.8 kb | ~455.6 kb | ~69.0 kb |
| **Physical Condensin spacing** |  |  | ~2.8 nm | ~4.3 nm | ~4.7 nm | ~13.0 nm | ~26.6 nm | ~4.0 nm |

Given the absence of periodic patterns or regular clusters along the chromatid axis, we assume for simplicity an equidistant spacing of Condensins along chromatid arms in order to predict the genomic spacing of the ∼1,400 Condensin holocomplexes on an average prometaphase HeLa chromatid. This calculation results in an average genomic distance between neighboring Condensins of ∼90 kb in prometa- and metaphase, which would be reduced to ∼70 kb by the binding of additional Condensin I complexes in anaphase (Table 1). If we further assume that each Condensin holocomplex defines a DNA loop, Condensins would be able to form ∼90 kb loops on prometaphase chromosomes, which would be spaced about 7 nm apart along the chromatid axis (123 Mb = 10.2 µm, 90 kb ≈ 7 nm), with most loops bound by Condensin I and only every fifth by Condensin II. This makes the clear prediction that Condensin II complexes should on average be spaced ∼35 nm apart along the chromatid axis.

Since STED super-resolution imaging has, in principle, a resolving power of ∼30 nm in 2D, we analyzed our STED data of prometaphase chromosomes for the limiting kleisin subunit of Condensin II (CAP-H2) to see if the number of detected localizations and their spatial distribution matched these predictions. Using 3D computational image analysis (Fig. S3D; for details see Materials and Methods), we could detect 16 spot-like localizations of Condensin II per µm of chromatid axis, slightly more than half of the 28 predicted from the abundance measurement by FCS-calibrated imaging. This indicates a combined labeling and detection efficiency of ∼60% by the combination of our primary and secondary antibody bearing the STED dye, making it very likely that most localizations represent single Condensin II complexes. We observed a median axial spacing of 56 nm (if projected onto the chromatid axis; Fig. S3D,G; for details see Materials and Methods). Assuming that we label and detect ∼60% of the complexes present, this number is in very good agreement with our predicted axial spacing of 35 nm from FCS-calibrated imaging. In anaphase the observed median axial spacing decreases further from 56 to 50 nm, while the distance from the axis increases from 115 to 133 nm, consistent with the average intensity and FWHM measurements (Fig. 3A,B).

### Data-driven three-step hierarchical loop model of mitotic chromosome compaction

Our integrated quantitative imaging data provide the absolute abundance, physical spacing and dynamic binding of Condensins on mitotic chromosomes. Together with the calibration of physical and genomic length of the mitotic genome in a human cell, these parameters allow us to estimate the maximum sizes of DNA loops that could be formed by Condensins to structure mitotic chromosomes. Taking into account the prevailing model of the loop extruding and stabilizing function of Condensin complexes (Alipour and Marko, 2012; Goloborodko et al., 2016A,B; Nasmyth, 2001), we therefore propose a three-step hierarchical loop model for how Condensins compact mitotic chromosomes (Fig. 4C). Condensin II is nuclear during interphase and its abundance on chromosomes is constant throughout mitosis (Fig. 1A,C). It likely promotes the initial axial compaction step in prophase and achieves this within the central 30-35% of the chromatid diameter (Fig. 3A,B). Based on the measured number as well as the measured physical and calculated genomic spacing of Condensin II complexes (Table 1), these loops would be large and about 450 kb in size if fully extruded. By contrast, the majority of Condensin I only gains access to mitotic chromosomes upon NEBD in prometaphase (Fig. 1A,C), when it binds to the central chromatid axis but also reaches more peripheral sites, occupying up to 50% of the chromatid diameter (Fig. 3A,B). The first phase of Condensin I binding leads to a 4- fold higher abundance compared to Condensin II on metaphase chromosomes (Fig. 1C,D) and likely drives lateral compaction in prometaphase and metaphase by subdividing the large Condensin II-mediated loops 4 times into sub-loops of a size of about 90 kb. Upon anaphase onset, a second wave of Condensin I binds to mitotic chromosomes (Fig. 1A,C), which then reaches a 5.6-fold higher abundance than Condensin II (Fig. 1C,D) and localizes even further away from the chromatid axis (Fig. 3A,B). These additional Condensin I complexes might drive the final axial shortening step by further subdividing the large Condensin II loops into 5-6 sub-loops, leading to a final average loop size of about 70 kb. The short (∼2 min) residence time of Condensin I as determined by FRAP (Fig. S2K) would allow such dynamic changes in sub-loop architecture to occur on mitotically relevant time scales.

Our proposed model for the Condensin-based formation and structural organization of mitotic chromosomes is based on the following parameters determined here and assumptions supported by the current literature. Condensin holocomplexes are defined by the number of limiting kleisin subunits which we determined by FCS; these are on average equidistantly spaced along the genome based on our super-resolution localizations; all Condensin holocomplexes define loop bases, with one Condensin per DNA loop as recently supported in the literature (Ganji et al., 2018). The endpoint states of each mitotic phase illustrated in our model thus represent the maximum size of fully extruded Condensin-mediated loops under these assumptions (Fig. 4C). Using this logical and quantitative framework thus allowed us to integrate our findings of two consecutive Condensin I recruitment waves (Fig. 1C), the higher abundance of chromosome-bound Condensin I (Fig. 1C,D) as well as the larger distance of Condensin I from the chromatid center (Fig. 3A,B) with the non-overlapping localization of the two Condensin complexes (Fig. 3C) and to propose a comprehensive model of Condensin II forming large prophase loops and Condensin I shortening these loops by the generation of smaller sub-loops in prometaphase and anaphase onset.

Since the direct visualization and size determination of single DNA loops in whole cells remains to our knowledge a technically unsolved challenge, our postulated loop arrangement based on our quantitative and super-resolved data is speculative and intended to generate testable hypotheses to further advance the field. We note that our imaging-based model for human cells is consistent with a biochemical study of drug-synchronized chicken cells published during the revision of this work, which proposed a conceptually similar arrangement of large Condensin II-based and smaller Condensin I-based loops (Gibcus et al., 2018). This orthogonal chromosome conformation capture (Hi-C) study furthermore computationally predicted a helical arrangement of Condensin II along the mitotic chromosome axis and provided a few diffraction-limited images of Condensin II localization of hypotonically spread chromosomes. Although our model did not predict a helical arrangement and it is not evident by manual inspection of our data (e.g. Fig. 3C), the Hi-C study prompted us to examine our extensive super-resolved data set of Condensin II localization from whole cells by unbiased computational image analysis and statistical data mining. This analysis did not detect significant helical or otherwise periodic patterns of Condensin II along the chromatid axis, which can therefore not be supported nor excluded by our data.

Importantly, our quantitative imaging data-driven model is consistent with data derived by orthogonal biochemical crosslinking approaches that have proposed similar loop sizes in mitotic human HeLa (Naumova et al., 2013) or HAP1 cells (Elbatsh et al., 2017 *Preprint*) as well as chicken cells (Gibcus et al., 2018). Our single cell imaging-based results, however, provide for the first time absolute quantitative parameters regarding the abundance, physical spacing relative to chromatid geometry as well as genomic spacing of both human Condensin complexes. This will be a valuable resource for future models of mitotic chromosome structure and compaction. One way to validate our model further in the future would be to directly resolve single Condensins on the individual loops of the DNA molecule of a mitotic chromatid. This will require new labeling and imaging methods for DNA and proteins that can provide isotropic nanometer scale resolution in all three dimensions as well as an imaging depth sufficient to penetrate deep into mitotic cells.

## Materials and methods

### Cell culture

HeLa Kyoto (HK) cells were obtained from S. Narumiya (Kyoto University) and grown in 1x high glucose DMEM (Thermo Fisher Scientific, cat.# 41965039) supplemented with 10% (v/v) FBS (Thermo Fisher Scientific; cat.# 10270106; qualified, E.U.-approved, South America origin), 100 U/ml penicillin-streptomycin (Thermo Fisher Scientific; cat.# 15140122), 2 mM L-glutamine (Thermo Fisher Scientific; cat.# 25030081) and 1 mM sodium pyruvate (Thermo Fisher Scientific; cat.# 11360070) at 37 °C and 5% CO_2_ in cell culture dishes (ThermoFisher) in a cell culture incubator. Cells were passaged every two to three days by trypsinization (Trypsin-EDTA (0.05%), phenol red (Thermo Fisher Scientific; cat.# 25300054)) at a confluency of 80-90%. Cells were confirmed to be mycoplasma-free on a regular basis.

### Cell line generation

HK cells were used for genome editing to express endogenously tagged proteins of interest (POI). For tagging SMC4 at the C-terminus and CAP-H at the N-terminus with mEGFP, zinc finger nucleases (ZFN) containing DNA binding sequences listed in Table S1 were purchased from Sigma Aldrich. The donor plasmids consist of the mEGFP cDNA sequence flanked by left and right homology arms listed in Table S2. ZFN pairs and the donor plasmid were transfected into HK cells as described in Koch et al., 2017 *Preprint*, and Mahen et al., 2014. For tagging CAP-D2 and CAP-D3 (Wutz et al., 2017) at the C-terminus, CAP-H and CAP-H2 at the C-terminus as well as CAP-H2 at the N-terminus (Cai et al., 2017 *Preprint*) with mEGFP, paired CRISPR/Cas9D10A nickases were used. The design of gRNAs was performed as described in Koch et al., 2017 *Preprint*, and Otsuka et al., 2016, (gRNAs listed in Table S1) and transfected together with donor plasmids (donor plasmids listed in Table S2) as described in Koch et al., 2017 *Preprint*, and Otsuka et al., 2016. For SMC4 and CAP-D2 only heterozygous clones were achieved after the first round of genome editing. Thus, after confirming by Sanger sequencing that the non-tagged alleles were not mutated, a second round of genome editing identical to the first round was performed on top of a heterozygous clone resulting in homozygous clones. The double endogenously tagged cell line HK CAP-H2-mEGFP CAP-H-Halo was generated by using the homozygous HK CAP-H2-mEGFP clone as parental cell line and performing genome editing based on the paired CRISPR/Cas9D10A nickase approach (gRNAs listed in Table S1; donor plasmid listed in Table S2). All genome-edited cell lines were confirmed to be mycoplasma-free on a regular basis. The tagging strategy was designed so that all Condensin isoforms could be tagged in the C-terminally mEGFP-tagged SMC4, CAP-D2, CAP-D3 and CAP-H cell lines. For CAP-H2 one theoretically predicted but not experimentally detected isoform could not be tagged in the C-terminally mEGFP-tagged cell line but all isoforms could be tagged in the N-terminally mEGFP-tagged cell line. FCS-calibrated imaging of metaphase cells did not reveal a significant difference between the abundance of CAP-H2 proteins on metaphase chromosomes between N- and C-terminally tagged clones (see Fig. S2F; two-sided Kolmogorov-Smirnov test, *p*-value = 0.1732, comparing data from 8 C-terminally tagged clones, *n* = 81 cells, and 5 N-terminally tagged clones, *n* = 41 cells). Thus, C-terminally mEGFP-tagged clones for all Condensin subunits were used throughout this study apart from Fig. S2F, which shows data from both C- and N-terminally tagged CAP-H and CAP-H2 clones, respectively.

### Cell line validation I: Junction PCR, SB, WB, confocal microscopy, mitotic timing analysis, Sanger sequencing

All single and double endogenously tagged cell lines were validated as described in Koch et al., 2017 *Preprint*, Mahen et al., 2014, and Otsuka et al., 2016 by performing Junction PCRs to test for integration and homozygosity (primers listed in Table S3), Southern Blot (SB) analysis with endogenous and GFP probes (and a Halo probe for double tagged cell lines) to test for homozygosity, local genome rearrangements and additional integrations of the GFP or Halo donors (probes listed in Table S5), Western Blot (WB) analysis to test for the expression of homozygously tagged proteins and absence of free GFP expression (antibodies listed in Table S4), confocal microscopy to test for correct localization of the tagged proteins, wide-field time-lapse microscopy and automated classification into mitotic phases (Held et al., 2010) to confirm that mitotic timing (from prophase to anaphase onset) was not perturbed, and Sanger sequencing. To confirm the correct localization of Halo-tagged CAP-H in the double endogenously tagged cell line, cells were stained with 100 nM HaloTag® TMR Ligand (Promega; cat.# G8251) in pre-warmed complete medium for 10 min at 37 °C and 5% CO_2_ and washed three times with pre-warmed 1x PBS (home-made), followed by changing to imaging medium (CO_2_-independent imaging medium (custom order from Thermo Fisher Scientific; based on cat.# 18045070; without phenol red); supplemented with 20% (v/v) FBS (Thermo Fisher Scientific; cat.# 10270106), 2 mM L-glutamine (Thermo Fisher Scientific; cat.# 25030081) and 1 mM sodium pyruvate (Thermo Fisher Scientific; cat.# 11360070)) and incubated for 30 min at 37 °C in the microscope incubation chamber prior to confocal microscopy as described in Koch et al., 2017 *Preprint*. In addition, co-immunoprecipitation (IP) against the GFP-tag was performed for all selected single homozygously mEGFP-tagged cell lines (HK 2x ZFN SMC4-mEGFP #82_68, HK 2x CRISPR CAP-D2-mEGFP #272_78, HK CRISPR CAP-D3-mEGFP #16, HK ZFN mEGFP-CAP-H #9, HK CRISPR CAP-H-mEGFP #86, HK CRISPR mEGFP-CAP-H2 #1, HK CRISPR CAP-H2-mEGFP #67, HK CRISPR CAP-H2-mEGFP CRISPR CAP-H-Halo) to confirm the assembly of Condensin subunits into pentameric complexes (antibodies listed in Table S4; for details see next paragraph).

### Cell line validation II: anti-GFP-IP and WB against Condensin subunits

GFP-Trap^®^_A beads (ChromoTek; cat.# gta-20) were used to pull down mEGFP-tagged proteins in HK cell extracts. Cells grown to 80-90% confluence were harvested either as an asynchronously cycling population or after 18 h incubation with 330 nM nocodazole (Noc; Merck; cat.# 48792) to yield a mitotically arrested cell population. Cell pellets were lysed in the following lysis buffer: 10% glycerol (Merck; cat.# 104093), 1 mM DTT (Sigma-Aldrich; cat.# D-9779), 0.15 mM EDTA (Merck; cat.# 324503), 0.5% Triton X-100 (Sigma-Aldrich; cat.# T8787), 1 tablet cOmplete EDTA protease inhibitor (Roche; cat.# 11873580001) and 1 tablet PhosSTOP (Roche; cat.# 04906837001). Lysis was performed on ice for 30 min and the suspension was centrifuged at 13,000 rpm and 4 °C for 10 min. The protein concentration of the cell extracts was determined by Bio-Rad Protein Assay (Bio-Rad; cat.# 500-0006) and 400 µg of total protein was used for IP. GFP-Trap^®^A beads were washed twice with 1x PBS (home-made) and used 1:1 diluted in PBS. 400 µg of total protein extracts were diluted 1:1 with PBS and incubated with the diluted GFP-Trap^®^A beads at 4 °C for 1 h. After the incubation, supernatant (SN) samples were taken and the beads were washed twice with lysis buffer diluted 1:1 with PBS. After two additional washing steps with PBS, 20 µl 4x NuPAGE™ LDS Sample Buffer (Thermo Fisher Scientific; cat.# NP0008) supplemented with 100 µM DTT was added to the beads and incubated at 65 °C for 5 min to elute the protein from the beads. Input (IN), SN and eluate (IP) samples were taken for WB against tagged and untagged subunits of the Condensin complexes (antibodies listed in Table S4).

### FCS-calibrated confocal time-lapse imaging (Sample preparation, FCS-calibrated confocal time-lapse imaging, data processing and analysis)

Samples for FCS-calibrated confocal time-lapse imaging were prepared as described in Cai et al., 2017 *Preprint*, Politi et al., 2017 *Preprint*. In brief, 2×10^4^ cells, which had been passaged the day before, were seeded into individual chambers of a Nunc™ 8-well LabTek™ Chambered Coverglass #1.0 (Thermo Fisher Scientific, cat.# 155411) and incubated overnight (ON) at 37 °C and 5% CO_2_ in a cell culture incubator. 2 h prior imaging, the medium was changed to imaging medium (see Cell line validation I) containing 50 nM SiR-DNA (Spirochrome; cat.# SC007; Lukinavičius et al., 2015) and 3.3 µM 500 kDa Dextran (Thermo Fisher Scientific; cat.# D7144) labelled with Dy481XL (Dyomics; cat.# 481XL-00; Dextran-Dy481XL produced in house).

FCS-calibrated confocal time-lapse imaging was performed as described in Cai et al., 2017 *Preprint,* and Politi et al., 2017 *Preprint* (Fig. S1D-H). FCS measurements of a 50 nM fluorescent dye solution of AlexaFluor488 (Thermo Fisher Scientific, cat.# A20000) in ddH_2_O were performed to estimate the confocal volume. FCS measurements in both the nucleus and the cytoplasm of wild-type cells not expressing mEGFP were performed to determine background fluorescence and background photon counts. FCS measurements in both the nucleus and the cytoplasm of wild-type interphase cells expressing free mEGFP as well as interphase cells homozygously expressing an mEGFP-tagged Condensin subunit were performed to estimate an experiment-specific calibration factor which was used to transform mEGFP fluorescence into mEGFP concentration (Fig. S1G). Prophase cells expressing an mEGFP-tagged Condensin subunit were automatically detected in low resolution imaging mode based on the DNA staining (SiR-DNA) using *CellCognition* (https://www.cellcognition-project.org/; Held et al., 2010). Upon recognition of a prophase cell, high-resolution time-lapse imaging of this cell through mitosis was triggered. Hereby, a *z*-stack covering the whole volume of the cell was acquired every 1.5 minutes for a total of 60 min in the mEGFP (mEGFP-tagged Condensin subunit), SiR-DNA (DNA), Dextran-Dy481XL (regions outside cells) and transmission channels. After each mitotic time-lapse one of the two daughter nuclei was detected and six FCS- measurements were acquired, three each inside and outside the nucleus, and used in addition to the interphase measurements for establishing the FCS calibration curve to transform mEGFP fluorescence intensities in each *z*-stack of a mitotic time-lapse video into protein concentrations. A 3D segmentation pipeline was used to detect and track cells through mitosis as well as to reconstruct chromosomal and cell surfaces from the SiR-DNA and Dextran-Dy481XL channels, respectively. (Cai et al., 2017 *Preprint*). To compensate for the variation of mitotic progression between individual cells, a mitotic standard time was computed based on geometric features of chromosomes and each cell was aligned to this mitotic standard time of 60 min (Cai et al., 2017 *Preprint*). FCS data processing, generation of calibrated images and data analysis to create cellular protein concentration and number maps (for details please refer to “Estimation of protein numbers from FCS-calibrated images”) including a multi-step quality control pipeline were performed as described in Cai et al., 2017 *Preprint*, Politi et al., 2017 *Preprint*, and Wachsmuth et al., 2015.

### Estimation of protein numbers from FCS-calibrated images

To estimate the number of proteins, fluorescent images of dividing cells were processed according to Cai et al., 2017 *Preprint*, and Politi et al., 2017 *Preprint*. Protein fluorescence intensities in image voxels were converted into number of proteins using the FCS calibration curve (Fig. S1G-H). 3D segmentations of the Dextran-DY481XL and SiR-DNA channels defined binary masks for the cell and chromatin/nuclear regions, respectively (Fig. S1I). The region in the cell but not in the chromatin region (XOR operation) was used to specify the cytoplasmic region. The total number of proteins was computed by adding up all proteins within the cell mask. Average concentrations in the chromatin/nucleus [*Chr*] and cytoplasm [*Cyt*] regions were computed by dividing the total number of proteins in the corresponding region by its volume. The FRAP results indicate that Condensin subunits can freely diffuse within the chromatin region (Fig. S2H-L). To compute the number of proteins bound to chromatin, we subtracted this freely diffusible protein pool. Between NEBD, mitotic standard stage 4, and the end of anaphase, mitotic standard stage 14 (Cai et al., 2017 *Preprint*), the nuclear envelope is not sealed. Thus, the concentration of the freely diffusible protein pool could be assumed to be equal to the measured cytoplasmic concentration: [*Chr*]*_bg_* = [*Cyt*]. This yielded the following corrected concentration and total number of chromatin-bound proteins

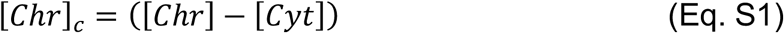

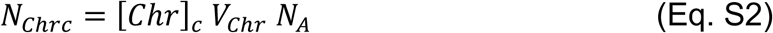

and a corrected total number of proteins in the cytoplasm

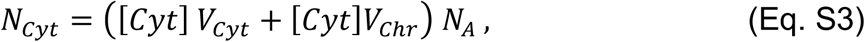

where *V_chr_* and *V_Cyt_* are the volumes of the chromatin region and cytoplasmic region, respectively, and *N_A_* the Avogadro constant. Before NEBD (prophase), we assumed that binding and unbinding of the Condensin subunits to chromatin is in equilibrium. The equilibrium constant *K_d_* was estimated from the concentrations in metaphase 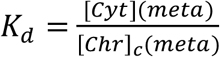. Thus, the freely diffusible nuclear concentration before NEBD is

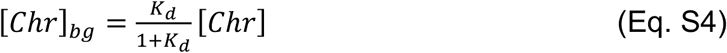

and the corrected chromatin-bound concentration

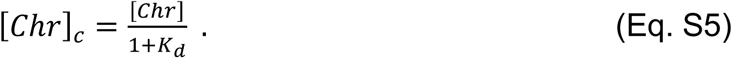

The number of proteins was calculated by multiplying these values with the chromatin volume. After nuclear envelope reformation, an equilibrium constant between chromatin-bound and free diffusing Condensin subunits could not be determined. Therefore we only computed the total number of proteins in the nucleus.

### FRAP experiments and analysis

Cells were seeded as for FCS-calibrated imaging. 2 h prior to imaging, the medium was changed to imaging medium (see Cell line validation I) containing 50 nM SiR-DNA to stain the DNA. Experiments were performed on a Zeiss LSM780 laser scanning microscope with an inverted Axio Observer (Zeiss, Oberkochen, Germany) operated by the ZEN2012 black software (Zeiss, Oberkochen, Germany). The microscope was equipped with a temperature-controlled incubation chamber (constructed in-house) and the temperature was set to 37 °C for imaging of living cells. Images were acquired using a C-Apochromat 40x/1.20 W Korr FCS M27 water immersion objective (Zeiss, Oberkochen, Germany) with an in-house built objective cap connected to an automated pump control system and spectral Gallium arsenide phosphide (GaAsP) detectors (Zeiss, Oberkochen, Germany). Cells in metaphase were manually selected based on their DNA staining and imaged every 20 s for 30-45 frames (9 *z* planes; *xyz* pixel size: 0.25 x 0.25 x 0.75 µm). Condensin-mEGFP was excited with 488 nm (Argon laser at 2%) and detected with GaAsP detectors at 500 - 535 nm. SiR-DNA was excited with 633 nm (HeNe laser at 0.3%) and detected with GaAsP detectors at 642 - 695 nm. After one pre-bleach image, a region of interest (ROI) covering approximately half of the chromatin (metaphase plate) in the 5th *z* plane was bleached using a squared region of 40×30 pixels (488 nm laser at 100%; 100 repetitions) and time-lapse movies were recorded.

A custom written ImageJ (Schindelin et al., 2012) script was used to extract the mean fluorescence intensity of the mEGFP-tagged POI and SiR-DNA in the bleached and unbleached chromatin region. To this end, the SiR-DNA channel was segmented to obtain the chromatin mask. The mask was then applied to the SiR-DNA and mEGFP channels previously filtered with an anisotropic 2D diffusion filter (Tschumperlé and Deriche, 2005). For every time point and *z* plane, the 1D profile of the SiR-DNA and mEGFP fluorescence intensity along the major 2D axis of chromatin was computed. Data close to the center of the 1D profile, which corresponds to the border of the bleaching ROI, was omitted to avoid boundary effects. Here, a gap of 14 pixels (3.5 µm) was used. The weighted average of the fluorescence intensity of the POI was computed using the SiR-DNA intensities as weights. The mean fluorescence intensity in the bleached (*F_b_*) and unbleached (*F_ub_*) was obtained as follows

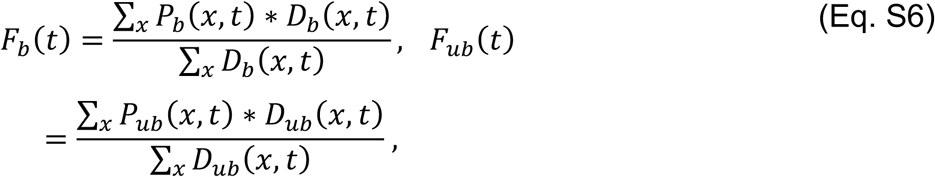

where *P_b,ub_*(*x,t*) and *D_b,ub_*(*x,t*) are the POI and SiR-DNA fluorescence intensities at position *x* along the DNA axis and at time *t*, respectively. The normalized difference between the unbleached and bleached region 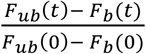 was used as readout for the residence time and immobile fraction (Gerlich et al., 2006).

Assuming a binding equilibrium of the POI with the chromatin and a constant total amount of protein, a system of differential equations for the unbleached and bleached region is obtained. The solution of the differential equations gives a closed expression for the normalized difference between unbleached and bleached regions:

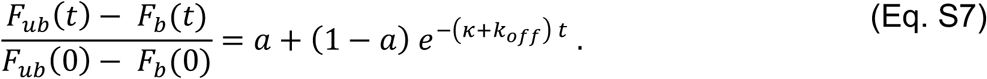

Here, the forward rate constant *κ* = *k_on_ B* is the product of the binding rate constant *k_on_* and the number of binding sites *B*, *k_off_* is the unbinding rate constant, and *a* is the immobile fraction. The benefit of using the difference between unbleached and bleached region is that all terms describing photo-bleaching cancel out and do not appear in Eq. S7. The residence time is given by

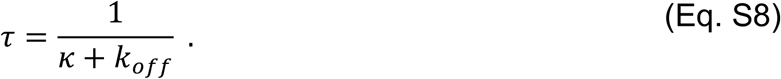

Different from classical point FRAP, where *τ* ≈ 1/*k_off_*, the residence time depends on the binding and unbinding rate constants. This is a consequence of the fact that a large portion of the proteins were bleached.

### Sample preparation for STED microscopy

To prepare HK cells homozygously expressing an mEGFP-tagged Condensin subunit for STED microscopy, 2.5×10^5^ cells, which had been passaged the day before, were seeded on squared glass cover slips (Marienfeld; cat.# 0107032), pre-washed with 1x PBS in Nunc™ 6-well plates (Thermo Fisher Scientific; cat.# 140685) and incubated ON at 37 °C and 5% CO_2_ in a cell culture incubator. On the next day the asynchronously cycling cell population was washed once with 1x PBS (+Ca^2+^/Mg^2+^; 0.9 mM CaCl_2_, 0.5 mM MgCl_2_; home-made). Pre-extraction was performed with 0.1% Triton™ X-100 (Sigma-Aldrich; cat.# T8787) in 1x PBS (+Ca^2+^/Mg^2+^) for 3 min at RT to permeabilize cell and nuclear membranes and remove soluble proteins, followed by two additional rounds of washing with 1x PBS (+Ca^2+^/Mg^2+^). All steps upto here were performed very carefully to not lose rounded-up and thus easily detachable mitotic cells. Cells were fixed with 4% PFA (Electron Microscopy Sciences; cat.# 15710) in 1x PBS (+Ca^2+^/Mg^2+^) for 15-20 min at RT. After three washing steps with 1x PBS, samples were quenched by incubation in 50 mM Tris-Cl (pH 7-8; home-made) for 5 min, followed by two washes with 1x PBS for 5 min each. Further permeabilization was carried out by incubation in 0.2% Triton™ X-100 in 1x PBS for 10 min, followed by three brief washing steps and two 5 min washes in 1x PBS.

Samples were blocked in 2% (w/v) BSA (Sigma-Aldrich; cat.# A2153) in 1x PBS for 1 h at RT. Incubation with the primary anti-GFP-antibody (AB; polyclonal rabbit anti-GFP-AB; MBL; cat.# 598; 1:500) in blocking solution (2% (w/v) BSA in 1x PBS) was performed ON at 4 °C. Three 5 min washes with blocking solution were followed by incubation with the secondary AB coupled to a STED dye (polyclonal goat anti-rabbit-Abberior STAR RED; Abberior; cat.# 2-0012-011-9; 1:500) in blocking solution for 1 h at RT. Samples were washed once with 1x PBS, and incubated with Hoechst 33342 (Thermo Fisher Scientific; cat.# H3570; 1:10,000) in 1x PBS for 10 min at RT, followed by three washes with 1x PBS for 5 min. Samples were mounted in DABCO-glycerol (1% (w/v) DABCO (Sigma-Aldrich; cat.# D27802), 90% (v/v) glycerol (Merck; cat.# 104093), in 1x PBS) on glass slides (Thermo Fisher Scientific; cat.# 12114682), sealed with Picodent twinsil™ (Picodent; cat.# 13001000) and kept at 4 °C in the dark until imaging. STED imaging was performed as soon as possible after sample preparation.

To prepare HK CAP-H2-mEGFP CAP-H-Halo cells for STED imaging, the same protocol was used apart from the following changes: The polyclonal primary ABs anti-GFP (chicken anti-GFP; Abcam; cat.# ab13970; 1:500) and anti-Halo (rabbit anti-Halo; Promega; cat.# G9281; 1:500) as well as the polyclonal goat secondary ABs anti-chicken-AlexaFluor594 (Thermo Fisher Scientific; cat.# A-11042; 1:500) and anti-rabbit-Abberior STAR RED were used in blocking solution. For primary AB staining, samples were pre-incubated with rabbit anti-Halo-AB for 1 h at RT prior to ON incubation with rabbit anti-Halo-AB and chicken anti-GFP-AB at 4 °C. The two secondary ABs were incubated together for 1 h at RT. No DNA staining with Hoechst was performed.

### STED imaging

STED imaging was performed on a combined Abberior STED and RESOLFT system (Expert line; Abberior Instruments, Göttingen, Germany) operated by the Imspector software (v0.13.11885; Abberior Instruments, Göttingen, Germany). Samples were imaged with an Olympus UPLSAPO 100x NA 1.4 oil immersion objective (UPLSAPO100XO/1.4; Olympus, Japan) on an Olympus IX83 stand (Olympus, Japan). The microscope was equipped with an incubation chamber (constructed in-house) and a temperature of 22.5 ± 0.2 °C was ensured by constant cooling to minimize sample drift and optimize optical performance. For STED imaging a 775 nm pulsed depletion laser (MPBC, Canada; 1.25 W total power, attenuated for actual imaging, AOTF set to 40%) was used which created a doughnut-shaped 2D depletion beam increasing the resolution laterally but not axially in comparison to a confocal microscope.

For diffraction-limited and STED imaging of mEGFP-tagged Condensin subunits stained with Abberior STAR RED, the dye was excited with a 640 nm pulsed excitation laser and emitted photons were recorded on an avalanche photodiode (APD) with a 650 - 720 nm bandpass filter in line sequential mode. For STED imaging the pulsed 640 nm excitation laser was triggered by the 775 nm pulsed STED depletion laser with 1 ns pulse length. A detection delay of 781.3 ps in respect to the excitation pulse was applied. In addition, DNA stained by Hoechst 33342 and the mEGFP-tagged Condensin subunit were imaged diffraction-limited in line sequential mode using 405 nm or 488 nm excitation lasers, respectively, and a 500 - 550 nm bandpass filter for detection on a separate APD. The pinhole was set to 1.2 airy units and the pixel dwell time to 2 µs.

For overview images of whole cells as shown in Fig. 2 a single *z* plane was acquired with the following settings: Eight line accumulations were performed, each of the following sequence: DNA (405 nm excitation laser, 10% (corresponding average power of 2.1 µW at the objective)) and GFP (488 nm excitation laser, 25% (28.8 µW)) channels were accumulated three times, whereas both the diffraction-limited (640 nm excitation laser, 2% (4.0 µW)) and the STED (640 nm excitation laser, 2% (4.0 µW); 775 nm depletion laser, 40% (166.3 mW)) channel of the immunostained Condensin subunit were accumulated five times, resulting in a total of 24 accumulations for diffraction-limited imaging of DNA and GFP as well as a total of 40 accumulations for diffraction-limited and STED imaging of the immunostained Condensin subunit. For acquiring *z*-stacks used for the analyses in Fig. 3A,B and Fig. S3, selected chromatids were imaged only in the DNA channel (diffraction-limited) and the immunostained Condensin subunit (STED) channel to minimize bleaching by using otherwise the same settings as described for overview images.

Mitotic cells in late prometaphase and early anaphase with chromatids having their longitudinal axes oriented parallel to the focus plane were selected for imaging based on their DNA and Condensin-mEGFP signals. The pixel size in *xy* was 20 nm. *Z*-stacks were acquired using the focus stabilizer/*z* drift control ZDC2 (Olympus, Japan; custom-modified for STED use) by focusing via the offset of the device and thereby ensuring precise axial positioning of each slice. To cover whole chromosomes in 3D, *z*-stacks of 29 slices and a *z* interval of 140 nm were acquired. By this *z* depth of about 4 µm it could be ensured that the whole 3D volume of not only straight but also slightly bent chromosomes was covered.

0.1 µm TetraSpeck™ Microspheres (Thermo Fisher Scientific; cat.# T7279) mounted on glass slides were imaged to align STED and excitation beams and to determine a potential offset between the diffraction-limited DNA and super-resolved immunostained Condensin-mEGFP channels. If necessary, an offset was corrected before performing the analyses shown in Fig. 3A,B and Fig. S3.

For two color STED microscopy of Condensin I and II as shown in Fig. 3C, chromatid regions of HK CAP-H2-mEGFP CAP-H-Halo cells stained as described above were imaged as follows: Eight line accumulations were performed, each of the following sequence: Both Condensin II (CAP-H2-mEGFP, AlexaFluor594; 594 nm excitation laser, 30% (3.6 µW)) and Condensin I (CAP-H-Halo, Abberior STAR RED; 640 nm excitation laser, 40% (95.6 µW)) channels were accumulated five times (resulting in a total of 40 accumulations) and imaged super-resolved (775 nm depletion laser, 40% (166.3 mW)) in line sequential mode using 605 - 625 nm and 650 - 720 nm band pass filters, respectively, for detection on separate APDs. The pinhole was set to 1.2 airy units and the pixel dwell time to 2 µs. The pixel size in *xy* was 20 nm. *Z*-stacks of 13 slices and a *z* interval of 140 nm were acquired.

Similar to the single color STED data, 0.1 µm TetraSpeck™ Microspheres mounted on glass slides were imaged to align STED and excitation beams and to determine a potential offset between the super-resolved immunostained Condensin I and Condensin II channels. If necessary, an offset was corrected before performing the co-localization analyses shown in Fig. 3C.

### Analysis of STED data: Single color STED image analysis – Condensin width and intensity distributions

ROIs containing straight or marginally bent chromatids mostly separable from neighboring chromatids were cropped for further analysis. A potential drift between consecutive *z* slices of the STED channel was determined by applying the MATLAB (MATLAB R2017a, The MathWorks, Inc., Nattick, Massachusetts, United States) *imregtform* function using rigid transformation and multimodal optimization. The transformation function representing the drift was applied to correct both the STED channel of the immunostained Condensin-mEGFP and the diffraction-limited DNA channel. A potential offset between the Condensin and DNA channels was determined based on *z*-stacks of 0.1 µm TetraSpeck™ Microspheres using the Channel alignment tool in the ZEN2012 black software (Zeiss, Oberkochen, Germany) and applied to the DNA channel to align Condensin and DNA channels. An in-house developed MATLAB script was used to manually segment the 3D Condensin volume from the STED channel by drawing the outline of the Condensin region on individual *z* slices. The corresponding chromatid volume was segmented similarly from the DNA channel as shown in Fig. S3A. Condensin and DNA *z*-stacks and their binary masks were interpolated along the *z* dimension to achieve an isotropic voxel size in *xy* and *z*. The segmented binary mask of DNA was extended by including any voxels that were not segmented as DNA but as Condensin. The interpolated Condensin *z*-stack was pre-processed using a 3D Gaussian filter (σ = 0.5).

All Condensin positive voxels within the interpolated binary mask were represented by their three orthogonal eigenvectors and associated eigenvalues where the largest eigenvector was used to cut the segmented Condensin volume at 100 nm spacing to generate a set of parallel cross sections orthogonal to the largest eigenvector. The centroids of each Condensin cross section were determined to adapt the direction of slicing based on the local vector generated from two neighboring centroids. Using these local vectors both Condensin and DNA volumes were cut again at 20 nm spacing to generate cross sections compatible with the local curvature of the Condensin volume as shown in Fig. S3A. The centroids of the final cross sections defined the Condensin axis (see Fig. S3D, yellow line). The length of Condensin axis per chromatid region was determined and used later for determining the number of Condensins per µm axis length. A small part of the Condensin volume from both ends of the chromatid was excluded from the analysis (see Fig. S3A, grey).

To calculate the width of Condensin cross sections within a sliding window of 2 µm length 2 μm, the cross sections were added up together and a one dimensional profile was generated by taking the sum projection of the intensities along *z* as shown in Fig. S3B. The width of Condensin was calculated by the full width at half maximum (FWHM) of the projected intensity profile shown in Fig. S3B. The sliding window was shifted by 200 nm to calculate the width of Condensin in a similar way. The mean of widths computed from all sliding windows per chromatid was taken as the Condensin width of this chromatid region. The chromatid width was calculated similarly. The Condensin width means of all chromatids per Condensin subunit and mitotic phase were combined to yield the means plotted in Fig. 3A. The chromatid width means of all chromatids for all Condensin subunits per mitotic phase were combined to yield means for prometaphase and anaphase. Since the chromatid width in anaphase was not significantly different from the chromatid in prometaphase, all chromatid width means independent of Condensin subunit and mitotic phase were pooled to yield the mean plotted in Fig. 3A.

To visualize the total intensity profile of Condensin subunits, all slices belonging to one chromatid region were added up to generate a 2D image (Fig. S3C, first panel), followed by sum projection. Profiles from all regions belonging to a particular Condensin subunit and mitotic phase were aligned to their maximum and merged to generate a combined profile. The profile was made symmetric by reflecting its left side to the right, taking the average and reflecting the average back to the left (see Fig. 3B, lower panels with total intensity profiles).

To generate the radial intensity profiles of Condensin subunits per mitotic phase, all cross sections belonging to one chromatid region were added up. Images from different regions belonging to a particular subunit and mitotic phase were combined by aligning them to a reference point at which the total intensity within a window size 0.4 and 0.6 times of the average chromatid width in *x* and *z*, respectively, reaches its maximum (Fig. S3C, second panel). An intensity profile through the reference point along the *x* axis (Fig. S3C, second panel) was taken (Fig. S3C, third panel). The left side of the profile was reflected to the right to compute the mean representing the radial profile (Fig. S3C, fifth panel). For intuitive visualization this profile was reconstructed into a 2D cross section (Fig. S3C, fourth panel) in which the intensity from the center to the periphery at any angle corresponds to the 1D profile. The mean chromatid width was used to exclude Condensin intensities outside the chromatid (Fig. S3C, white region in fifth panel).

### Analysis of STED data: Double colour STED image analysis – co-localization analysis of Condensin I and II

Cropping of ROIs containing chromatid arms, drift correction between consecutive *z*-slices and correction of shift between the Condensin I and Condensin II channels were performed similarly to the single color STED data. An initial manual segmentation of the combined Condensin volume was performed only in the Condensin I channel and the same binary mask was applied to segment the Condensin II volume (see Fig. 3D, first two panels). Co-localization analysis was performed by only considering the detected high density voxels of Condensin I and Condensin II by further segmenting the initial combined Condensin volume. A global threshold for each Condensin was automatically detected by applying the Otsu method (Otsu, 1979). High density voxels of Condensin signals were segmented by adapting the MATLAB Bradley local thresholding function (MATLAB Central, File Exchange; https://nl.mathworks.com/matlabcentral/fileexchange/40854-bradley-local-image-thresholding), whereby the global threshold was combined with a local threshold for a window of size 9×9 to reduce the amount of over-/under-segmentation. The percentage of non-overlapping voxels was calculated by the number of non-overlapping voxels divided by the geometric mean of the number of Condensin I and Condensin II positive voxels multiplied by 100. Condensin I and Condensin II positive voxels were combined to determine the intensity correlation between Condensin I and II using normalized cross correlation.

### Analysis of STED data: Single colour STED image analysis – Automatic spots detection, clustering, counting, centroid determination as well as coordinate-based distance measurements

Intensity peaks in each slice of a Condensin STED *z*-stack were localized with FIJI (Schindelin et al., 2012) using the ThunderSTORM v1.3 plugin (https://github.com/zitmen/thunderstorm; Ovesný et al., 2014) with the following camera settings: readoutnoise = 0.0, offset = 0.0, quantumefficiency = 1.0, isemgain = false, photons2adu = 1.0, pixelsize = 20.0 nm). Background noise was removed with a B-spline wavelet filter with order of 2 and scale of 4, followed by an approximation of peak position using a local maximum threshold of 3*SD (Wave.F1). The sub-pixel positions of the intensity peaks were found with the following settings: estimator = [PSF: Integrated Gaussian], sigma = 2.2, fitradius = 2, method = [Maximum likelihood], full_image_fitting = false, mfaenabled = false).

For each region the detected *x*, *y*, and *z* positions within the manually segmented Condensin region (spots) were clustered using the DBSCAN (Density-based Spatial Clustering of Applications with Noise) v1.1-1 algorithm (Ester et al., 1996) in RStudio v3.4.2 (RStudio Team (2015); Boston, Massachusetts, United States) with an eps value of 45 and minPts set to two. The *z* values were scaled by a factor of 1/8 to account for the at least 8 times poorer resolution in *z* (diffraction-limited) in comparison to *xy* (STED super-resolved) due to the anisotropy of 2D STED and to make peaks adjacent in *z* more likely to be placed in the same cluster. For the least abundant Condensin II subunit CAP-H2 most clusters were well separated from each other and assumed to represent one Condensin subunit and, thus, one Condensin complex. However, to account for the possibility of several Condensin subunits being present in the same high intensity cluster, the cluster number per micrometer was corrected by not allowing clusters to be larger than 3000 arbitrary intensity units and, thus, using 3000 arbitrary intensity units or multiples thereof as threshold for counting two or more Condensin subunits per cluster. The number of clusters per micrometer Condensin axis length was calculated from the Condensin axis length per chromatid region (determined in: Analysis of STED data: Single color STED image analysis – Condensin width and intensity distributions) and the intensity-corrected number of clusters. The mean number of clusters representing the number of CAP-H2 subunits for all respective chromatid regions in prometaphase and anaphase was determined by averaging among all CAP-H2 chromatid regions per mitotic phase.

The performance of the outlined procedure of automatic spots counting, clustering and intensity-correction to account for multiple proteins per cluster was confirmed by manual spots counting of three chromatid regions of CAP-H2 in early anaphase using IMARIS (v9.0.1; Bitplane, Zurich, Switzerland), demonstrating a maximum deviation of the automated spots counting of 18% from manual counting.

Cluster centroids were determined and used for the following distance measurements: Pairwise distances between all cluster centroids were calculated using MATLAB *IPDM* function (MATLAB Central, File Exchange; https://nl.mathworks.com/matlabcentral/fileexchange/18937-ipdm-inter-point-distance-matrix) and the minimum distance of neighboring clusters was taken as the 2D nearest neighbor (NN) distance (Fig. S3D,E). To calculate the distance between cluster centroids and the Condensin axis (CA), a distance transformed image of the Condensin axis was generated in which the intensity of each pixel represents its distance from the axis. Thus, the coordinates of the cluster centroids were used to determine their CA distances (Fig. S3D,F). Cluster centroids were projected orthogonally on the Condensin axis to obtain a 1D representation of the distribution. The distance between neighboring 1D projected cluster centroid pairs on the Condensin axis was calculated and used as readout for the axial spacing (AS) of Condensins (Fig. S3D,G).

### Airyscan imaging of live SMC4-mEGFP cells and determination of whole cell chromatid length

2.5×10^4^ HK SMC4-mEGFP cells, which had been passaged the day before, were seeded into individual chambers of a Nunc™ 8-well LabTek™ II Chambered Coverglass #1.5 (Thermo Fisher Scientific, cat.# 155409) and incubated ON at 37 °C and 5% CO_2_ in a cell culture incubator. 2 h prior to imaging, the medium was changed to imaging medium (see Cell line validation I) containing 50 nM SiR-DNA to stain the DNA and cells were kept at 37 °C until imaging.

Experiments were performed on a Zeiss LSM880 laser scanning microscope with an inverted Axio Observer (Zeiss, Oberkochen, Germany) operated by the ZEN 2.3 black software (Zeiss, Oberkochen, Germany) with AiryscanFast imaging option. The microscope was equipped with a temperature-controlled incubation chamber (constructed in-house). Although live cells were imaged, the temperature was set to RT to slow down cell division and reduce cellular movement during the acquisition of individual *z*-stacks. Images were acquired using a C-Apochromat 40x/1.20 W Korr FCS M27 water immersion objective (Zeiss, Oberkochen, Germany). Cells in late prometaphase, metaphase or early anaphase were manually selected based on their DNA and Condensin signal. SMC4- mEGFP was excited with 488 nm (Argon laser at 3.5%) and SiR-DNA was excited with 633 nm (HeNe laser at 2.8%). Detection was performed on the Airy disc consisting of 32 GaAsP-PMT (photomultiplier tube) detectors by using a combined 495 - 550 nm BP (band-pass) and 570 nm LP (low-pass) filter. AiryscanFast imaging mode was used. The *xy* pixel size was 50 nm. A *z*-stack with a *z* interval of 200 nm of a manually defined size was acquired to cover the complete DNA volume of a cell. Airyscan *z*-stacks were Airyscan processed in 3D using the ZEN 2.3 black software and automatic settings.

The total chromosome length of a mitotic cell was computed by using the length of a small pre-segmented Condensin volume and the ratio of the total intensity of Condensin signal within the entire chromosomal volume and the total intensity of the segmented Condensin volume. To this end, the SMC4-mEGFP channel was used to quantify the Condensin intensity and the SiR-DNA channel was used to segment the chromosomal volume. The shift between the two channels was determined for individual cells by cross correlation within a 350 nm x 350 nm x 600 nm neighborhood and applying the average offset among all cells to the DNA channel of all cells. A custom-written MATLAB script was used to manually segment a small Condensin volume largely distinguishable from other regions (Fig. 4A). Individual *z* slices were pre-processed using a 2D Gaussian filter (σ = 2.5 for DNA, σ = 2.0 for Condensin). DNA and Condensin signals were segmented by combining local and global thresholds as described previously (Hériché et al. 2014). The segmented slice containing the maximum number of Condensin positive voxels was dilated using diamond shape structuring element of radius 9 pixels and eroded using diamond shape structuring element of radius 7 pixels to extract the rim for quantifying Condensin background intensity. A histogram of the rim intensities was created and the average background intensity was calculated from the histogram without considering the lowest 10% and the highest 50% voxels.

The segmented DNA volume was applied to the Condensin signal and the manually segmented Condensin volume to exclude Condensin voxels outside the chromosomal volume. The manually segmented Condensin region was interpolated to yield isotropic voxel sizes in *xy* and *z*. Boundary voxels were detected by morphological operation and their pairwise distances were calculated using the MATLAB *IPDM*. The pair of voxels with the largest distance was selected to determine the vector for slicing orthogonal cross sections of the manually segmented Condensin volume at 50 nm spacing. The centroids of each cross section were determined to define a central axis of the segmented Condensin volume. The length of the central axis was computed and considered as the length of the segmented Condensin volume. The total chromosome length of each cell was calculated from the total intensity of the Condensin signal within the entire DNA volume divided by the total intensity of the manually segmented Condensin volume multiplied by its Condensin axis length. 3-5 Condensin regions per cell were manually segmented and used for determining the total chromatid length per cell. For each cell the mean based on the calculations from different segmented Condensin regions was determined. For each mitotic phase, the mean of total cellular chromatid length was determined by averaging among three cells.

### Statistical methods

The relative precision of FCS-calibrated confocal imaging was determined by comparing the quantification of proteins on chromatin for cells in metaphase. The intra-assay variability given by the coefficient of variation (CV; CV = standard-deviation/mean) was ∼11%. This error accounts for the biological variability and precision of the imaging and image analysis pipeline. The inter-assay variability, computed by calculating the mean and standard deviation between all experiments, yielded a CV of 19.6%. This CV also accounts for the variability generated by the FCS-calibration method.

For the FCS-calibrated imaging data and the FRAP data in Fig. 1 and Fig. S2 means, medians, standard deviations and interquartile ranges were computed using *R* (https://www.r-project.org/) and *ggplot*. Robust linear regression (*rlm* function in the *R* package *MASS*) was used to compute the FCS calibration curve for each experiment (e.g. Fig. S1G). To fit the exponential model to the FRAP data (Fig. S2 J-L) the Levenberg-Marquardt algorithm was used as implemented in the function *nlsLM* from the *R* package *minpack.lm*. Parameter distributions were computed by bootstrap (Efron and Tibshirani, 1994). To this end, a random sample containing the same number of cells as in the original data set was created by random sampling with replacement. For each sampled set the exponential model was fitted to the data. This operation was repeated *N* times (here *N* = 300) yielding parameter distributions. From the parameter distributions medians and interquartile ranges were computed. To compare the amount of CAP-H2 on metaphase chromosomes the two-sided Kolmogorov-Smirnov test was used as implemented in the *R* function *ks.test*.

For the STED data in Fig. 3 means and standard deviations were calculated using MATLAB (MATLAB R2017a, The MathWorks, Inc., Nattick, Massachusetts, United States). Significance tests of Condensin widths (Fig. 3A) were based on Student’s two-tailed t-tests and performed using Microsoft Excel 2007 (Microsoft Corporation; Redmond, Washington, United States). The following thresholds for *p*-values were used to indicate significance: non-significant *p*≥0.05, * *p*<0.05, ** *p*<0.01, *** p<0.001.

Medians for nearest neighbor distance (NN), distance of cluster centroids from the central Condensin axis (CA) and axial spacing (AS) of CAP-H2 based on the coordinates of detected and clustered spots in the CAP-H2 STED data (Fig. S3E-G) were calculated using MATLAB (MATLAB R2017a, The MathWorks, Inc., Nattick, Massachusetts, United States).

Means and standard deviations for whole cell chromatid lengths (Fig. 4B) were calculated using Microsoft Excel 2007 (Microsoft Corporation; Redmond, Washington, United States).

### Online supplemental material

Figure S1 shows the validation of genome-edited cell lines expressing homozygous mEGFP-tagged Condensin subunits and the FCS-calibrated imaging methods. Figure S2 shows the absolute quantification of Condensin-mEGFP subunits over mitosis by FCS-calibrated imaging and the results from FRAP experiments of Condensin-mEGFP subunits on the metaphase plate. Figure S3 illustrates the methods to determine the width (FWHM) and intensity profiles of Condensin subunits from *z*-stacks of chromatid regions acquired in 2D STED mode as well as the methods for automatic spot detection and clustering of CAP-H2 STED data as well as distance measurements based on cluster centroids. Table S1 lists gRNA and ZFN sequences and Table S2 donor plasmids used for genome editing. Table S3 lists primers for Junction PCR. Table S4 lists antibodies used for WB. Table S5 lists probes used for SB. The source code for the image and data analysis methods can be found online. *MitoSys* contains and links to the source code for analyzing FCS-calibrated confocal time-lapse data through mitosis and generating time-resolved cellular protein concentration and number maps (Fig. 1B-D, Fig. S2A-E). *Segmentation_single_cell* contains source code used to segment chromosomes in metaphase cells (Fig. S2F). *FRAP* contains the source code for analyzing FRAP data (Fig. S2I-L). *Chromatid_structure_analysis* contains the source code for the analysis of single color (Fig. 3A,B) and double color (Fig. 3C) Condensin STED data, the distribution analysis of Condensin spots (Fig. S3E-G) as well as the determination of whole cell chromatid length (Fig. 4A,B). *zStackSpotPicker* contains the source code for detecting and clustering Condensin spots (Fig. S3D). The latest version of the source code can also be found at https://git.embl.de/grp-ellenberg/condensin_map_walther_jcb_2018.

## Acknowledgements

We are grateful to Yin Cai and Malte Wachsmuth for initially setting up the FCS-calibrated 4D imaging pipeline. We thank the Flow Cytometry Core Facility (FCCF) at EMBL for support in cell sorting of genome-edited cells, as well as the Advanced Light Microscopy Facility (ALMF; especially Christian Tischer and Aliaksandr Halavatyi, Zeiss and Abberior Instruments (especially Christian Wurm and Tim Grotjohann) for support in the initial set up of STED imaging conditions and help with the STED microscope. We gratefully acknowledge Arina Rybina for help with setting up imaging conditions at the Zeiss LSM880 AiryscanFast microscope, Kira Elaine Detrois for manual spots counting and segmentations, Jean-Karim Hériché for advice on statistical analyses, Carl Barton and Jeremy Tempkin for help with scripts, and the EMBL design team for graphics support. We thank Christian Häring, Daniel Gerlich, Sara Cuylen-Häring, Stephanie Alexander, Merle Hantsche-Grininger and Bianca Nijmeijer for critically reading the manuscript. We are grateful to Christian Häring and members of the Häring lab, Daniel Gerlich, Jan-Michael Peters, Jonas Ries, Edward Lemke, Michael Knop, and all current and former members of the Ellenberg lab, especially M. Julia Roberti, Wanqing Xiang, Yin Cai and Merle Hantsche-Grininger, for fruitful discussions. This work was supported by grants to J.E. from the EU-FP7- SystemsMicroscopy NoE (Grant Agreement 258068), the EU-FP7-MitoSys (Grant Agreement 241548) and the 4D Nucleome/4DN NIH Common Fund (U01 EB021223) as well as by EMBL (N.W., M.J.H., A.Z.P., B.K., M.K., Ø.Ø., M.L., J.E.). N.W. and Ø.Ø. were supported by the EMBL International PhD Programme (EIPP). The authors declare no competing financial interests.

## Author contributions

N.W. and J.E. conceived the project, designed the experiments and wrote the manuscript. N.W. performed all experiments. B.K. and M.K. generated and N.W., B.K. and M.K. validated genome-edited cell lines. A.Z.P. and N.W. analyzed FCS-calibrated images. A.Z.P. developed the FRAP analysis pipeline and performed analyses together with N.W. N.W. and M.L. set up STED imaging conditions. M.J.H. developed computational pipelines for STED analysis and the determination of whole cell chromatid length and performed analyses together with N.W. Ø.Ø. and N.W. set up the pipeline for automated spot detection and clustering and M.J.H. performed analyses together with N.W.

## Abbreviations

AB: antibody
APD: avalanche photodiode
CV: coefficient of variation
FCS: fluorescence correlation spectroscopy
FWHM: full width at half maximum
GaAsP: Gallium arsenide phosphide
HK: HeLa Kyoto
IN: input
IP: immunoprecipitation / eluate
NEBD: nuclear envelope breakdown
Noc: nocodazole
ON: overnight
POI: protein of interest
ROI: region of interest
SB: Southern Blot
SMC: structural maintenance of chromosomes
SN: supernatant
STED: stimulated emission depletion
WB: Western Blot
ZFN: zinc finger nuclease

