## Supplementary Materials for "A quantitative map of human Condensins provides new insights into mitotic chromosome architecture"

*Supplemental figure legends and table titles*

**Fig. S1: Validation of genome-edited homozygous mEGFP knock-in cell lines and methods for FCS-calibrated confocal imaging and determination of protein numbers on mitotic chromosomes.**

**(A-C)** Genome-edited HeLa Kyoto cells homozygously mEGFP-tagged for Condensin subunits were validated **(A)** for homozygosity and absence of both genomic rearrangements and extra integrations of the mEGFP donor by Southern Blot (SB), **(B)** anti-GFP-immunoprecipitation (IP) of asynchronous (Noc -) and mitotically arrested cells (Noc +) and Western Blot (WB) against the other Condensin subunits to ensure the incorporation of tagged subunits into functional pentameric Condensin complexes, homozygous expression of the mEGFP-tagged protein and absence of the expression of free GFP, as well as **(C)** non-perturbance of mitotic timing (prophase to anaphase onset; mean + SD) by wide-field time-lapse microscopy and automatic classification into mitotic phases. For (A-C) WT: wild-type. In (A) the expected sizes for mEGFP-tagged and non-tagged SMC4 by the GFP and endogenous probes used for SB are 3.5 kb and 2.8 kb, respectively. For (B) Noc: nocodazole, IN: input, SN: supernatant, IP: eluate. In (C) one representative experiment out of three independent experiments is shown (WT: $n$ = 87, SMC4-mEGFP: $n$ = 71). Validation data in (A-C) is exemplarily shown for the homozygous HK SMC4-mEGFP cell line. All other homozygous Condensin-mEGFP cell lines used in this work were validated similarly. **(D-I)** Methods for FCS-calibrated confocal imaging and determination of protein numbers on mitotic chromosomes (details for Fig. 1C,D). **(D)** FCS-measurements are acquired in the nucleus (point 1) and cytoplasm (point 2) of cells expressing the protein of interest (POI) tagged with mEGFP (e.g. SMC4-mEGFP), mEGFP alone and wild-type cells (WT) that do not express mEGFP. At each FCS position the fluorescence intensity (*FI*) is measured in a 9x9 region of interest (ROI). HeLa Kyoto interphase cell expressing SMC4-mEGFP and stained with SiR-DNA. The nuclear border is indicated with an orange dashed line. Scale bar: 10 µm. **(E)** Photon counts at the positions indicated in (D) are recorded over time. **(F)** Autocorrelation curves are computed (dots) and fitted to a two-component diffusion model (solid lines) to extract the number of molecules $N_1$ and $N_2$ in the focal volume. The concentration $C$ is calculated by dividing the number of molecules with the effective focal volume $V_{eff}$ and the Avogadro constant $N_A$. The effective focal volume has been determined from a measurement of the diffusion time of an Alexa488 dye solution in the same experiment. (D-F) are representative data from 414 FCS measurements from 140 SMC4-mEGFP cells (2-6 FCS points per cell) and 5 experiments. On average, 426 FCS measurements from 141 cells were acquired per genome-edited cell line in 5 independent experiments. **(G)** The FCS-calibration curve obtained from measurements of cells expressing the POI-mEGFP (SMC4-mEGFP, $n$ = 118 measurements from 36 cells), mEGFP alone ($n$ = 19 measurements from 12 cells) or WT cells ($n$ = 10 measurements from 5 cells) for background (BG) correction. Concentrations are plotted against the fluorescence intensities and fitted to a linear function (straight line indicated). Representative data from one experiment is shown. **(H)** The pixel fluorescence intensity ($FI_p$, left panel) is converted to concentrations (middle panel) by applying the linear relationship from (G) and protein numbers (right panel) by multiplying with the voxel volume $\Delta x\ \Delta y\ \Delta z$ and the Avogadro constant $N_A$. A SMC4-mEGFP metaphase cell is shown. Scale bar: 10 µm. **(I)**

Exemplary segmentation of cell boundary (left panel) and DNA (middle panel) based on extracellular fluorescently labelled Dextran or SiR-DNA staining, respectively, of the SMC4-mEGFP cell shown in (H). A transmission image as well as the 3D reconstructed cell (grey) and chromatin (magenta) volumes are shown in the right panel. Scale bar: 10 µm.

**<u>Fig. S2:</u> Quantitative live cell data of Condensin subunits in mitotic cells and FRAP of Condensin subunits on the metaphase plate.**

**(A-G)** Quantitative live cell data of HeLa Kyoto cells with homozygously mEGFP-tagged Condensin subunits (details for FCS-calibrated imaging/Fig. 1): SMC4 (Condensin I/II, grey), CAP-D2 (dark magenta) and CAP-H (light magenta, Condensin I) as well as CAP-D3 (dark green) and CAP-H2 (light green, Condensin II). **(A)** Total protein numbers as function of the mitotic standard time. **(B)** Number of proteins in the cytoplasm. This is computed by summing up all proteins in the cytoplasmic region. After NEBD to the end of anaphase Eq. S3 (see Materials and Methods) is used. **(C)** Concentration of proteins bound to chromosomes (left panel) or in the nucleus (right panel). Before NEBD Eq. S5 is used, after NEBD up to the end of anaphase Eq. S1 is used. **(D)** Concentration of proteins in the cytoplasm. (A-D) Mean (colored lines) and SD (light grey areas) are shown (~20 cells per subunit (range: 10-36) with 3-7 independent experiments; 36, 22, 10, 17 and 17 cells from 7, 3, 5, 3, and 4 experiments for CAP-H, CAP-H2, CAP-D2, CAP-D3 and SMC4, respectively). **(E)** Fraction of proteins bound to chromatin measured in metaphase (Fig. 1C,D). The median, upper and lower quartiles as well as 1.5 times the interquartile range are shown (~20 cells per subunit (range: 10-36) with 3-7 independent experiments; 36, 22, 10, 17 and 17 cells from 7, 3, 5, 3, and 4 experiments for CAP-H, CAP-H2, CAP-D2, CAP-D3 and SMC4, respectively). **(F)** Fold difference in the Condensin I subunits bound to chromatin between CAP-H and CAP-D2 for different homozygous clones. For CAP-H both C- and N-terminally mEGFP-tagged clones are shown. The clones used in this study are indicated in bold. The number of cells is 22, 7, 6, 5, 19, 2 for CAP-H N_z9, N_z30, N_z133, C_c70, C_c86, N_z156, respectively, and 26 for CAP-D2 C_c272c78. **(G)** Fold difference in the Condensin II subunits bound to chromatin between CAP-H2 and CAP-D3 for different homozygous clones. For CAP-H2 both C- and N-terminally mEGFP-tagged clones are shown. The clones used in this study are indicated in bold. The number of cells is 7, 9, 24, 7, 10, 10, 14, 7, 6, 7, 8, 6, 7 for CAP-H2 C_c11, N_c176, N_c1, C_c37, N_c30, N_c27, C_c67, C_c19, N_c76, N_c28, N_c159, C_c36, N_c123, respectively, and 18, 8, 6 for CAP-D3 C_c16, C_c48, C_c57, respectively. **(H-L)** FRAP experiments of Condensin subunits on the metaphase plate. **(H)** FRAP of mEGFP-tagged Condensin subunits (green) was performed by bleaching half of the metaphase plate (chromosomes stained by SiR-DNA) at time point 0 as indicated in the pre-bleach image ROI (yellow square) and acquiring a time-course ($\Delta t$ = 20 s). Maximum projected images of *z* planes 3-7 are shown. A Gaussian blur (σ=1) was applied to the images for presentation purposes. Scale bar: 10 µm. The images are representative for *n* = 9 (CAP-H) and *n* = 15 (CAP-H2) cells from 2 and 3 independent experiments, respectively. **(I)** Normalized fluorescence intensity difference in the unbleached ($F_{ub}$) and bleached ($F_b$) chromatin region. Shown is mean ± SD of *n* = 16 (SMC4, grey), *n* = 14 (CAP-D2, dark magenta), *n* = 9 (CAP-H, light magenta), *n* = 15 (CAP-D3, dark green) and $n$ = 15 (CAP-H2, light green) cells. 2-4 independent experiments per Condensin subunit were acquired. **(J)** The mean of fluorescence recovery traces (points) and fits to Eq. S7 (solid line) are shown for each subunit. **(K)** The residence time $\tau$ is defined by Eq. S8 and plotted for the different Condensin cell lines (median, upper and lower quartile as well as 1.5 times the interquartile range). **(L)** The immobile fraction is defined by Eq. S7 and plotted for the different cell lines (median, upper and lower quartile as well as 1.5 times the interquartile range). The data plotted in (J-L) is based on 2-4 independent experiments with $n$ = 16 (SMC4, grey), $n$ = 14 (CAP-D2, dark magenta), $n$ = 9 (CAP-H, light magenta), $n$ = 15 (CAP-D3, dark green) and $n$ = 15 (CAP-H2, light green) cells. The statistics were computed by bootstrapping the data (for details see Materials and Methods).

**<u>Fig. S3:</u> Analysis of Condensin STED data to determine the sub-chromosomal distribution of Condensin subunits and automatic spot detection for distance measurements of the Condensin II subunit CAP-H2.**

**(A-C)** Sub-chromosomal distribution of Condensin subunits based on 3D STED data of Condensin subunits in HeLa Kyoto cells. **(A)** $z$-stacks of immunostained mEGFP-tagged Condensin subunits were acquired super-resolved in 2D STED mode. Chromosomes stained by Hoechst were acquired diffraction-limited. Left panel: Condensin and chromatid were segmented manually slice by slice. A representative $z$ plane of a CAP-D3 chromatid in anaphase is shown. Scale bars: 500 nm. Right panel: 3D visualization of Condensin (green) in the segmented DNA volume (magenta). Condensin and DNA channels were interpolated along $z$ to achieve an isotropic voxel size in $x$, $y$ and $z$. The central Condensin axis was determined and cross sections along the chromatid diameter and perpendicular to the Condensin axis compatible with the local curvature of the Condensin volume were selected. Zoom-ins of cross sections and the corresponding normalized intensities of the Condensin and DNA channels are provided. Scale bars: 500 nm. **(B)** To calculate the width of Condensin, cross sections within a sliding window of 2 µm length were added up together and a 1D profile was generated by taking the sum projection of the intensities along $z$. The width of Condensin was calculated by the full width at half maximum (FWHM) of the projected intensity profile. The sliding window was shifted by 200 nm repeatedly, the Condensin width was calculated in a similar way for all 2 µm sliding windows and the Condensin width of the chromatid region was taken as the average among all sliding windows. The Condensin width of one particular subunit and mitotic phase was calculated by taking the mean from all chromatid regions belonging to this particular subunit and mitotic phase. The chromatid width was calculated similarly. **(C)** To visualize the total Condensin intensity profile per chromatid region, all cross sections belonging to one Condensin region were added up to generate a combined 2D image (first panel). Combined images calculated from different regions belonging to a particular subunit and mitotic phase were combined by aligning them to a reference point. An intensity profile through the reference point along the $x$-axis (second panel) is plotted (third panel). The left side of the profile was reflected to the right to compute the mean intensity profile representing a 1D radial profile. For intuitive visualization this profile was reconstructed into a 2D cross section in which the intensity from the center to the periphery at any angle corresponds to the 1D profile (fourth panel). The mean chromosome width was used to exclude the profile outside the chromosome marked in white (fifth panel; see radial intensity profiles in the upper panels of Fig. 3B). In (A) and (B) the same CAP-D3 chromatid in anaphase representative for *n* = 20 chromatids is shown. In (C) average data from *n* = 20 CAP-D3 chromatids in anaphase is shown. **(D-G)** Automatic spot detection and clustering for distance measurements of the Condensin II subunit CAP-H2. **(D)** Spots in *z*-stacks of CAP-H2 2D STED data were detected automatically and clustered (upper panel; different colors indicate different clusters). The cluster centroids (magenta) were determined and their nearest neighbor distances (NN, orange) as well as distances from the yellow central Condensin axis (CA, blue) were calculated in 2D (middle panel). Cluster centroids were projected onto the central Condensin axis and their 1D distance representing their axial spacing (AS, green) was calculated (lower panel). A representative prometaphase chromatid of CAP-H2 is shown with overview images on the left and zoom-ins (turquoise boxes) on the right. Scale bar overview: 500 nm. Scale bar zoom-in: 200 nm. **(E)** Histograms of the nearest neighbor distance (NN) of cluster centroids of CAP-H2 are plotted for prometaphase (left) and anaphase (right). The median is plotted as an orange line (dashed for prometaphase, continuous for anaphase) and the exact value is indicated. **(F)** Histograms of the distance of cluster centroids of CAP-H2 from the central Condensin axis (CA) are plotted for prometaphase (left) and anaphase (right). The median is plotted as a blue line (dashed for prometaphase, continuous for anaphase) and the exact value is indicated. **(G)** Histograms of axial spacing (AS) of cluster centroids of CAP-H2 projected in 1D onto the central Condensin axis are plotted for prometaphase (left) and anaphase (right). The median is plotted as a green line (dashed for prometaphase, continuous for anaphase) and the exact value is indicated. Prometaphase: *n* = 12; anaphase: *n* = 16).

**<u>Table S1:</u> gRNA and ZFN binding sequences for genome editing.**

**<u>Table S2:</u> Donor plasmids for genome editing.**

**<u>Table S3:</u> Primers for Junction PCR.**

**<u>Table S4:</u> Antibodies for WB.**

**<u>Table S5:</u> Probes for SB.**

Fig. S1

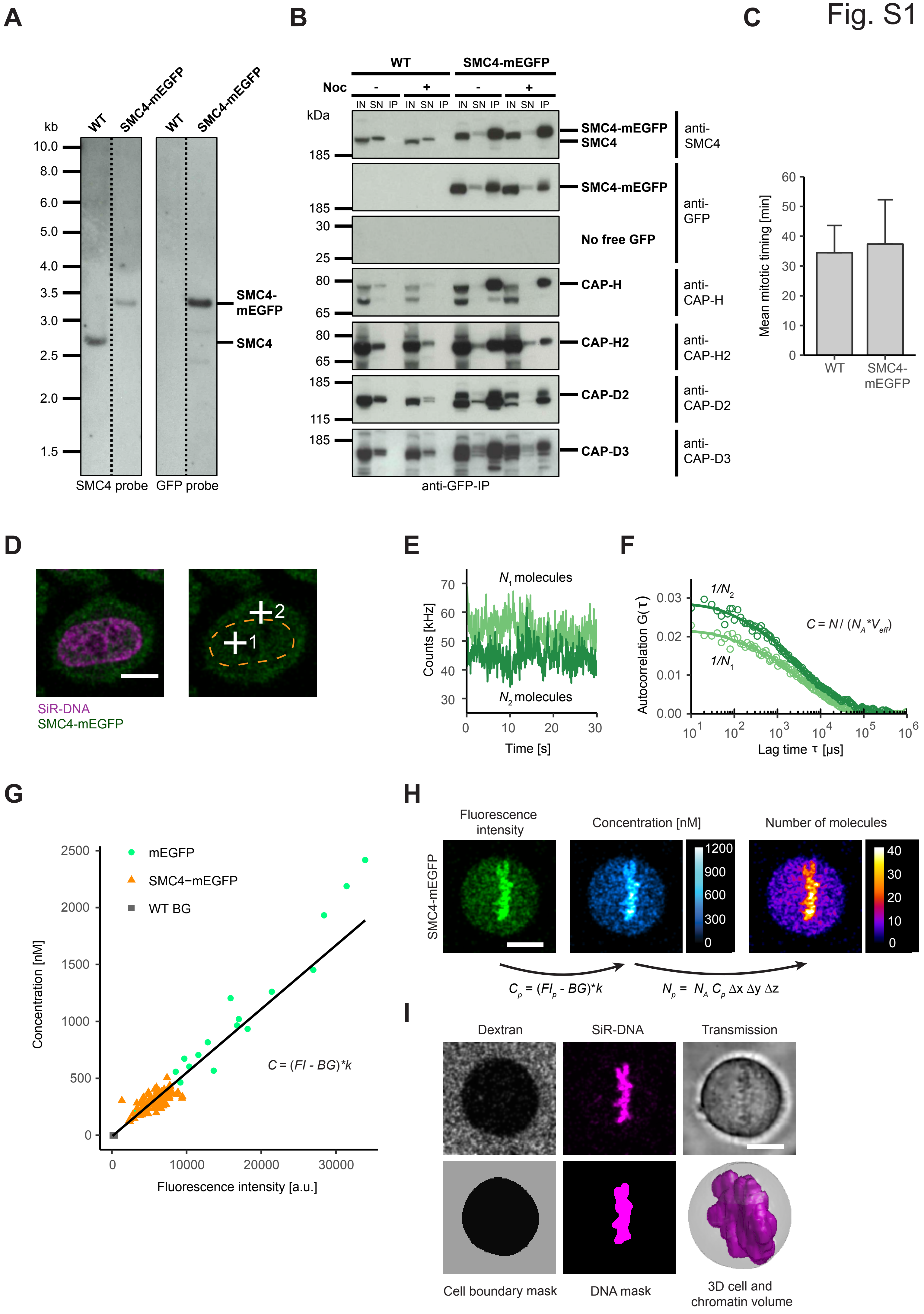

A
kb
WT
SMC4-mEGFP
WT
SMC4-mEGFP
10.0
8.0
6.0
5.0
4.0
3.5
3.0
2.5
2.0
1.5
SMC4-mEGFP
SMC4
SMC4 probe
GFP probe
B
WT
SMC4-mEGFP
Noc
-
+
-
+
IN SN IP IN SN IP IN SN IP IN SN IP
kDa
185
185
30
25
80
65
80
65
185
115
185
SMC4-mEGFP
SMC4
anti-SMC4
SMC4-mEGFP
anti-GFP
No free GFP
CAP-H
anti-CAP-H
CAP-H2
anti-CAP-H2
CAP-D2
anti-CAP-D2
CAP-D3
anti-CAP-D3
anti-GFP-IP
C
Mean mitotic timing [min]
60
50
40
30
20
10
0
WT
SMC4-mEGFP
D
2
1
SiR-DNA
SMC4-mEGFP
E
$N_1$ molecules
$N_2$ molecules
Counts [kHz]
70
60
50
40
30
0
10
20
30
Time [s]
F
Autocorrelation G(τ)
0.03
0.02
0.01
0.00
$1/N_2$
$1/N_1$
$C = N / (N_A * V_{eff})$
$10^1$ $10^2$ $10^3$ $10^4$ $10^5$ $10^6$
Lag time τ [μs]
G
mEGFP
SMC4−mEGFP
WT BG
Concentration [nM]
2500
2000
1500
1000
500
0
$C = (FI - BG)*k$
0
10000
20000
30000
Fluorescence intensity [a.u.]
H
SMC4-mEGFP
Fluorescence intensity
Concentration [nM]
1200
900
600
300
0
Number of molecules
40
30
20
10
0
$C_p = (FI_p - BG)*k$
$N_p = N_A C_p \Delta x \Delta y \Delta z$
I
Dextran
SiR-DNA
Transmission
Cell boundary mask
DNA mask
3D cell and chromatin volume

Fig. S2

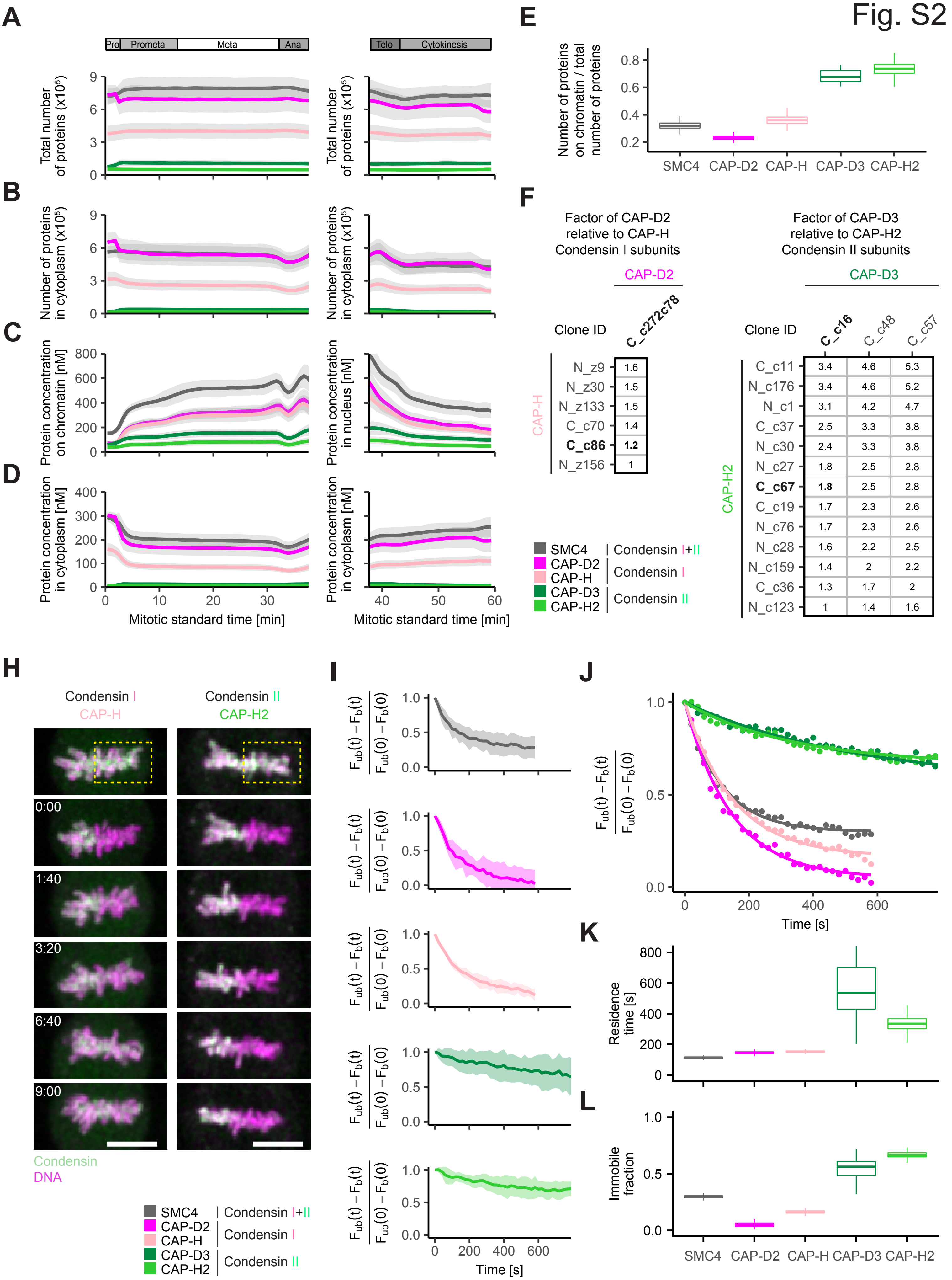
A
Pro Prometa Meta Ana
Telo Cytokinesis
Total number of proteins (x10^5)
B
Number of proteins in cytoplasm (x10^5)
C
Protein concentration on chromatin [nM]
Protein concentration in nucleus [nM]
D
Protein concentration in cytoplasm [nM]
Mitotic standard time [min]
E
Number of proteins on chromatin / total number of proteins
SMC4 CAP-D2 CAP-H CAP-D3 CAP-H2
F
Factor of CAP-D2 relative to CAP-H Condensin I subunits
CAP-D2
Clone ID
C_c272c78
CAP-H
N_z9 1.6
N_z30 1.5
N_z133 1.5
C_c70 1.4
C_c86 1.2
N_z156 1
Factor of CAP-D3 relative to CAP-H2 Condensin II subunits
CAP-D3
Clone ID C_c16 C_c48 C_c57
CAP-H2
C_c11 3.4 4.6 5.3
N_c176 3.4 4.6 5.2
N_c1 3.1 4.2 4.7
C_c37 2.5 3.3 3.8
N_c30 2.4 3.3 3.8
N_c27 1.8 2.5 2.8
C_c67 1.8 2.5 2.8
C_c19 1.7 2.3 2.6
N_c76 1.7 2.3 2.6
N_c28 1.6 2.2 2.5
N_c159 1.4 2 2.2
C_c36 1.3 1.7 2
N_c123 1 1.4 1.6
SMC4 | Condensin I+II
CAP-D2
CAP-H | Condensin I
CAP-D3
CAP-H2 | Condensin II
H
Condensin I
CAP-H
Condensin II
CAP-H2
0:00
1:40
3:20
6:40
9:00
Condensin
DNA
SMC4 | Condensin I+II
CAP-D2
CAP-H | Condensin I
CAP-D3
CAP-H2 | Condensin II
I
(F_ub(t) − F_b(t)) / (F_ub(0) − F_b(0))
Time [s]
J
Time [s]
K
Residence time [s]
L
Immobile fraction
SMC4 CAP-D2 CAP-H CAP-D3 CAP-H2

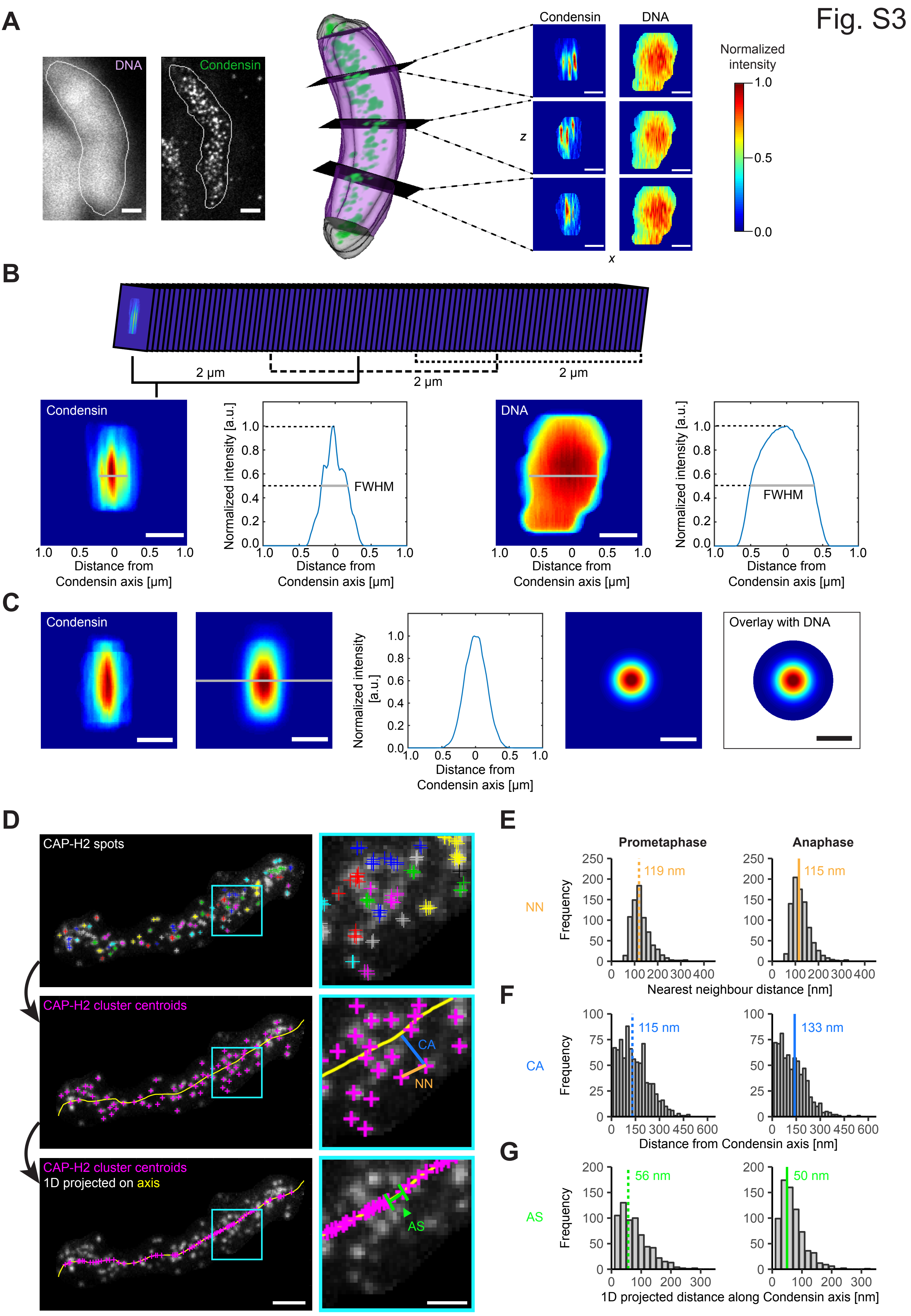

Fig. S3
A
DNA
Condensin
Condensin
DNA
z
x
Normalized intensity
1.0
0.5
0.0
B
2 µm
2 µm
2 µm
Condensin
Normalized intensity [a.u.]
FWHM
DNA
Distance from Condensin axis [µm]
C
Condensin
Overlay with DNA
D
CAP-H2 spots
CAP-H2 cluster centroids
CA
NN
CAP-H2 cluster centroids
1D projected on axis
AS
E
Prometaphase
Anaphase
NN
Frequency
119 nm
115 nm
Nearest neighbour distance [nm]
F
CA
115 nm
133 nm
Distance from Condensin axis [nm]
G
AS
56 nm
50 nm
1D projected distance along Condensin axis [nm]

**Table S1: gRNA and ZFN binding sequences for genome editing.**

| Cell line | ZFN binding sequences (5'-3') with cut site in lower case | Sequence of antisense gRNA (5‘-3‘) | Sequence of sense gRNA (5‘-3‘) |
|---|---|---|---|
| HK ZFN SMC4-mEGFP | TACATACTCCCTAaactagATCATGAAACTGGTT | | |
| HK CRISPR CAP-D2-mEGFP | | AGGAAGTCTGTTCCTGTCCT | GGGTATCCTGTAGGGTGACC |
| HK CRISPR CAP-D3-mEGFP | | GGTGGGAGGCGCTGTTTAGT | AGCGCCTCCCACCAGTGTCC |
| HK ZFN mEGFP-CAP-H | CCTCCCGGCCCAGGTgagccGGGCGGTCGGGAGGCGCG | | |
| HK CRISPR CAP-H-mEGFP | | AGGGAAGACCAAAACATGCA | CAAGGAGATTGAGTTCACTA |
| HK CRISPR mEGFP-CAP-H2 | | ACCGCAGGGCTGCCTTCCGA | CCGTTCCCTCCCGGACATGG |
| HK CRISPR CAP-H2-mEGFP | | TGCCTCGGTGCTCCCCACTC | GACCTACGCTGCCCCCTCCA |
| HK CRISPR CAP-H2-mEGFP CRISPR CAP-H-Halo | | AGGGAAGACCAAAACATGCA | CAAGGAGATTGAGTTCACTA |

**Table S2: Donor plasmids for genome editing.**

| Donor plasmid | ENSEMBL transcript ID* | Sequence 5' homology arm* | Sequence of 3' homology arm* |
|---|---|---|---|
| pSMC4-donor-Cterm-mEGFP | ENST00000357388.7 | Chromosome 3: 160433007-160433806 | Chromosome 3: 160433810-160434710 |
| pNCAPD2-donor-Cterm-mEGFP | ENST00000315579.9 | Chromosome 12: 6531413-6532212 | Chromosome 12: 6531413-6532212 |
| pNCAPD3-donor-Cterm-mEGFP | ENST00000281189.4 | Chromosome 11:134152947-134153746 | Chromosome 11: 134152144-134152943 |
| pNCAPH-donor-Nterm-mEGFP | ENST00000240423.8 | Chromosome 2: 96335033-96335829 | Chromosome 2: 96335874-96336655 |
| pNCAPH-donor-Cterm-mEGFP | ENST00000240423.8 | Chromosome 2: 96372549-96373348 | Chromosome 2: 96373352-96374151 |
| pNCAPH2-donor-Nterm-mEGFP | ENST00000299821.15 | Chromsome 22: 50507538-50508337 | Chromosome 22: 50508341-50509140 |
| pNCAPH2-donor-Cterm-mEGFP | ENST00000299821.15 | Chromosome 22: 50522573-50523372 | Chromosome 22: 50523376-50524175 |
| pNCAPH-donor-Cterm-Halo | ENST00000240423.8 | Chromosome 2: 96372549-96373348 | Chromosome 2: 96373352-96374151 |

*Ensembl release 90 (August 2017)

**Table S3: Primers for Junction PCR.**

| Gene | Forward primer (sequence 5‘-3‘) | Reverse primer (sequence 5‘-3‘) |
|---|---|---|
| SMC4 C-term | GTATGTGCTCCTATGTATGCATCCTCC | AGACTTCTAATACAATACCGAAAGTCCTTCC |
| NCPAD2 C-term | GCCTTCTCACCTTGGCTTCTTAGTAATGTG | CTCTTCCCACAAATGCTGGTAGCATTCTTC |
| NCAPD3 C-term | GAAGGATTTCCACCTCCACGCTCTG | TGTGTCCGACTCCAAAGGTTATGCTC |
| NCAPH N-term | CCCTGGCCTTTACTCCCATAATTCAATC | CTTCCTAGTGTTCTCACCTGTTCCCATG |
| NCAPH C-term | GGAAAGGTCTCATAGCTGCTTAGTCCTGAG | CCTGTATGACCTATCCAGGTAGCACATAGC |
| NCAPH2 N-term | GATACCTCCATCCCTAGGTCAGAGTCTG | AGCTGCAGATCTGGTAGTTGTCCTAATACAG |
| NCAPH2 C-term | GATGACTTTCTAGAGCCTGAGGAGTACATGG | CTTGTTTCCAGGAGCATCAGATCCATGC |
| EGFP | TCAAGGAGGACGGCAACATC | GATGTTGCCGTCCTCCTTGA |
| Halo | TCGAAGAATACATGGACTGGCTGCACC | GGTCTGGTTTGTCGGATTTGCCCATAC |

**Table S4: Antibodies for WB.**

| Name of AB | Dilution | Raised in | Supplier | Catalogue number |
|---|---|---|---|---|
| anti-SMC4 | 1:1000 | rabbit | Bethyl Laboratories | A300-064A |
| anti-CAP-D2 | 1:1000 | rabbit | Bethyl Laboratories | A300-601A |
| anti-CAP-D3 | 1:1000 | rabbit | Bethyl Laboratories | A300-604A |
| anti-CAP-H | 1:3000 | rabbit | Bethyl Laboratories | A300-603A |
| anti-CAP-H2 | 1:1000 | rabbit | Bethyl Laboratories | A302-275A |
| anti-SMC2 | 1:1000 | rabbit | Bethyl Laboratories | A300-056A |
| anti-CAP-G | 1:1000 | rabbit | Bethyl Laboratories | A300-602A |
| anti-CAP-G2 | 1:1000 | rabbit | Bethyl Laboratories | A300-605A |
| anti-GFP | 1:1000 | mouse | Roche | 11814460001 |
| anti-Halo | 1:1000 | mouse | Promega | G9211 |

**Table S5: Probes for SB.**

| Gene | Probe (sequence 5‘-3‘) |
|---|---|
| SMC4 C-term | AATTTGCTTGCATGCTATTTATGCTTGCAGTTTACAGGGGCAGACAGCCAGTATTTACAAAATATTTGTCTAGTTCATCTTTTCTTAAATTTTCAAGGCATTTTTAACATGAGGATTGAGAAATTGTATTTGATAAGGAGAAAGAATAGCAAGAAGTATAATACTTAGCTTGCTCTTTAAATCTTAAAACTTTGGTATGTGCTCCTATGTATGCATCCTCC |
| NCPAD2 C-term | AGTCACCTTCCCGTCCTCCTTATCCCCAGCTGGGTTGCAACCAAATTGCCAGAGTGACCTAAGACCAGATCTTTGTCTCCAGTTCTTTTTTTATTACTCCAAAAACACAACCAAAGCAGCATCTCATCCAATTCTTGTTTGTTTGTTTTTAATAGTTTTTATTTTTCAGAGCAGTTTTAGGTTCAAAGCAAAATTGAGCAGAAAGTACAGGGAGTTCCCTTCTAC |
| NCAPD3 C-term | TCAAGCTATGCATCAAGCTTGGTACCGAGCTCGGATCCACTAGTAACGGCCGCCAGTGTGCTGGAATTCGCCCTTCTTCCCTGGTGGATGGGTGATTGATAGCATTTTTAGTGTGCATGCCTAACCACTGAAATGCTAGGAAGAAGAGACAAACCCAACGTAAGAGCAAATAACAAAGAATGAAGCAATGTTTGGGTTTTTTTGAAGCTTTTATAAATGAGGTTAAACAAATGCCAAATTACGGTGTGTTAGTACAAGTCTGTAGCATCTGTGTACAGCAACATCCATCCAAAGGCAGAATCACAGCCTCGTATCGGCCTCGAGTCTTGAGTCACATTCATATCTTAAAATACACGAGTATTCCTTTTCCGTCTAGTCTCAGTTATAGGCAGCTACTTCCCAGACTCTACTTTCACTTCTCTCTACTTT |
| NCAPH N-term | ATAGGAAGTGTCTTTGAGCTCAGTAACCATCGGTTGAATGAATGGATGCATAATATCAAAGGCAAGGAGAGAAACCACGGAAAGAAGATTTGGAGAATTAAAGCATGCCTCTAATTCCAAGTTTAGTTTCTCTCTTAGACAAGGTTCT |
| NCAPH C-term | AGTAACAGAACTACTTGGATCAACATCTATAATATTGTTCCAAATTGCAAGCAAAGTTAATTCCCCTATTCCTCTGAGGTTAGGGGAAGAACTTGTACATTTGTGTGTGTGCTGCATTCTGTGGAGATGGACAAATCTGAGCACTGCTCTC |
| NCAPH2 N-term | GCCTAGGTCATCTTCTGATGCATTTACCCTGAAGTGTCATTTATTTCTATGATTTTAAATATTGCCTGTATTAGGACAACTACCAGATCTGCAGCTCCATCCTAGATCTTTCCTCCAAATCTACTTTACATATCCAGCTGCTCTCTTGACATCTAGTGGGTGTCACGCAAACATCTCATTTAACATATTTAAAGTGGAACTCTTCATTTTTCTGCTTAAACCCCAGCTTTTCCCATCTCGGAAAATGACACCGCTGTTCAACCAAAAGCAAAAGCTAAATGTGGCTATAGTGAAGCCCCATGCTTAACACTCAGTTCATCCATG |
| NCAPH2 C-term | TTGCTGACGGAAAGCATTCCAAGTGCATGCCTTGCCTGAACTAACCACGTTATCTATTTGCAATAAACCCATTTCTTAAAAGGAGGTGGGTAACTCTCT |
| EGFP | CACATGAAGCAGCACGACTTCTTCAAGTCCGCCATGCCCGAAGGCTACGTCCAGGAGCGCACCATCTTCTTCAAGGACGACGGCAACTACAAGACCCGCGCCGAGGTGAAGTTCGAGGGCGACACCCTGGTGAACCGCATCGAGCTGAAGGGCATCGACTTCAAGGAGGACGGCAACATCCTGGGGCACAAGCTGGAGTACAACTACAACAGCCACAACGTCTATATCATGGCCGACAAGCAGAAGAACGGCATCAAGGTGAACTTCAAGATCCGCCACAACATCGAGGACGGCAGCGTGCAGCTCGCCGACCACTACCAGCAGAACACCC |
| Halo | TGCATTGCTCCAGACCTGATCGGTATGGGCAAATCCGACAAACCAGACCTGGGTTATTTCTTCGACGACCACGTCCGCTTCATGGATGCCTTCATCGAAGC |
